# Antibody immunogenicity prediction and optimization with ImmunoSeq

**DOI:** 10.1101/2025.08.14.670305

**Authors:** Qiaojing Huang, Yi He, Kai Liu

## Abstract

Therapeutic antibody development faces persistent immunogenicity challenges from anti-drug antibodies (ADA). Identifying peptide fragments presented by major histocompatibility complex is the central challenge in predicting immunogenicity. Here, we presented ImmunoSeq, an interpretable and applicable method for immunogenicity prediction. ImmunoSeq addresses this by deploying complementary k-mer (k=8-12) peptide libraries: a positive library of immunologically safe peptides from fragmented human proteins/antibodies, and a negative library of murine antibody fragments capturing evolutionary-selected immunogenic triggers. For candidate antibodies, we generate all possible k-mer peptides and compute hit rate by summing positive hits (+1.0) and negative hits (-0.2 penalty) normalized against total peptide number. Higher hit rate predicts lower ADA risk, with residue-level resolution enabling precise localization of immunogenic hotspots. ImmunoSeq demonstrated superior ADA correlation and humanness classification accuracy compared to deep learning models, while accurately predicts ADA reductions in humanization, enabling sufficient sequence optimization for humanness. By leveraging dual-library discrimination principles of self/non-self-peptide, ImmunoSeq provides a robust, interpretable solution for immunogenicity prediction and sequence optimization.

## Main

Immunogenicity, the tendency of therapeutic macromolecules to induce anti-drug antibodies (ADA), remains a critical bottleneck in biotherapeutic development^1–5^. ADA can neutralize drug activity, diminish efficacy, or trigger adverse reactions, including hypersensitivity, autoimmune disorders, and even life-threatening cytokine release syndrome, thereby jeopardizing patient safety and undermining clinical success^6–7^. With the rapidly expanding pipeline of therapeutic biologics (e.g., monoclonal antibodies, fusion proteins, and gene therapies), early prediction and mitigation of immunogenicity have become imperative to reduce attrition and optimize therapeutic outcomes^8–13^. A central challenge in immunogenicity prediction lies in identifying T cell epitopes: short peptide fragments (typically 8-12 amino acids) derived from the drug, which are presented by major histocompatibility complex (MHC) molecules on antigen-presenting cells to activate T cells and drive ADA formation^14–17^. However, the MHC system is highly polymorphic (with thousands of alleles), and T cell receptor (TCR) recognition is highly context-dependent, which is shaped by the structural conformation of peptide-MHC complexes and surrounding microenvironmental cues, making it difficult to accurately predict which peptides will trigger an immune response. This uncertainty has limited the reliability of existing approaches, leaving a critical gap in rational immunogenicity mitigation.

Current immunogenicity prediction methods fall into distinct categories, each with notable limitations. MHC binding predictors (e.g., NetMHCpan^18–19^, MHCflurry^20–21^, BigMHC^22^) estimate peptide-MHC affinity but focus narrowly on binding, neglecting downstream processes such as antigen processing or TCR recognition. T cell epitope models (e.g., PanPep^23^, pMTnet^24^, NetTCR^25–27^, NeoRanking^28^, TEIM^29^, catELMo^30^) integrate MHC binding with TCR interaction features using machine learning (ML) models or protein language models, but rely heavily on limited annotated epitope data, leading to poor generalization for novel sequences. Sequence homology-based approaches^31–34^ (e.g., BLAST-based alignment, germline homology scoring, Human String Content^35^, T20 scores^36^) compare drug sequences to human proteins to infer “self-similarity” but lack granular resolution, as even small non-self-peptide regions can drive robust immunogenicity. Recent machine learning and deep learning (DL) models^37–39^ (e.g., DeepImmuno^40^, Hu-mAb^41^, Sapiens^42^, AbNatiV^43^) improve performance by leveraging language models to learn sequence patterns of human proteins and antibodies, aiming to assess antibody humanness, but remain black boxes, offering little insight into which specific residues drive risk and thus limiting their utility for sequence optimization.

To address these limitations, we present ImmunoSeq, a novel, interpretable approach rooted in the biological principle of immune tolerance: the human immune system is tolerant to self-proteins, which do not elicit ADA. ImmunoSeq constructs a self-peptide library by systematically fragmenting >20,000 human protein sequences and >1 million healthy human antibody sequences from the OAS^44–46^ paired dataset into k-mer peptides (k=8-12, matching dominant MHC-presented lengths), thereby generating a virtual repository of millions of immunologically safe peptides. Complemented by a negative non-self-peptide library derived from >80,000 mouse antibody sequences also from OAS paired dataset, this approach incorporates evolutionary selected immunogenic peptides for penalty calibration. For candidate therapeutic antibodies, we similarly fragment their sequences into k-mer peptides, quantifying both positive hits (matches against the self-peptide library) and negative hits (matches against the non-self-peptide library), assigning + 1.0 to each positive hit and -0.2 to each negative hit. These accumulated values of all hits, normalized against the total peptide number, yield the global hit rate. Theoretically, a higher hit rate indicates greater similarity to self-proteins, thus predicting lower ADA risk. Critically, ImmunoSeq achieves residue-level resolution of immunogenic contributions by quantifying per-residue hit rate - calculated as the sum of assigned values across all k-mer peptides containing the residue, normalized against the total peptide number. Residues with low hit rate are flagged as potential immunogenic hotspots, enabling precise localization of risky regions and guiding iterative mutation design to enhance the overall hit rate.

Validated on 217 therapeutic antibodies, ImmunoSeq correlates well with ADA incidence and accurately predicts immunogenicity changes during humanization. ImmunoSeq also exhibited comparable accuracy to DL methods in classifying human sequences across different species. Furthermore, ImmunoSeq shows superior practical utility in iterative sequence optimization: when generating mutation candidates to boost overall hit rate, model-recommended mutation sets aligned closely with experimental observations in 25 humanized antibody pairs. Specially, the top-ranked mutations (e.g., top 5 of recommendations) were highly concordant with experimentally validated beneficial mutations. This top enrichment, where the most impactful mutations are prioritized, enables efficient, targeted optimization, outperforming ML/DL methods in interpretability and actionable guidance. By grounding predictions in the biological principles of immune tolerance and self-similarity, ImmunoSeq bridges the gap between immunogenicity risk assessment and rational sequence engineering.

## Results

### ImmunoSeq Workflow

ImmunoSeq translates immune tolerance principles, where evolutionary pressures select antibodies capable of discriminating self from non-self, into a predictive computational framework through dual-library peptide matching (Fig. 1a). The framework quantitatively evaluates therapeutic antibody sequences against two context-specific peptide repertoires: a positive self-peptide library integrating >20,000 human proteome sequences and >1 million paired human antibody sequences from OAS database, capturing germline and affinity-matured sequences that have undergone negative selection; and a negative non-self-peptide library comprising >80,000 paired mouse antibody sequences from OAS database serving as archetypal immunogenic triggers due to their ∼30% sequence divergence from human equivalents. Critical to biological relevance, all sequences (including input antibody sequence and library sequence) are fragmented into 8-mer to 12-mer short peptides corresponding precisely to MHC class I (8-10) and class II (10-12) major binding cleft dimensions, ensuring coverage of T-cell receptor engagement zones critical for immunogenicity. To score the hit rate, the simplest way is to match the input peptide fragment against the library, which assigns +1.0 for self-peptide library hits (tolerance signatures) and -0.2 for non-self-peptide library hits (immunogenic penalties) to reconcile library size disparities (∼10^6^ vs. ∼10^5^) while maximizing clinical relevance. The global hit rate is thus computed by fragmenting antibody heavy and light chains into overlapping peptides, calculating chain-normalized hit rate, and averaging values to eliminate length confounders, finally yielding a metric between -0.2 (high immunogenicity) and 1 (high tolerance).

**Fig. 1.**
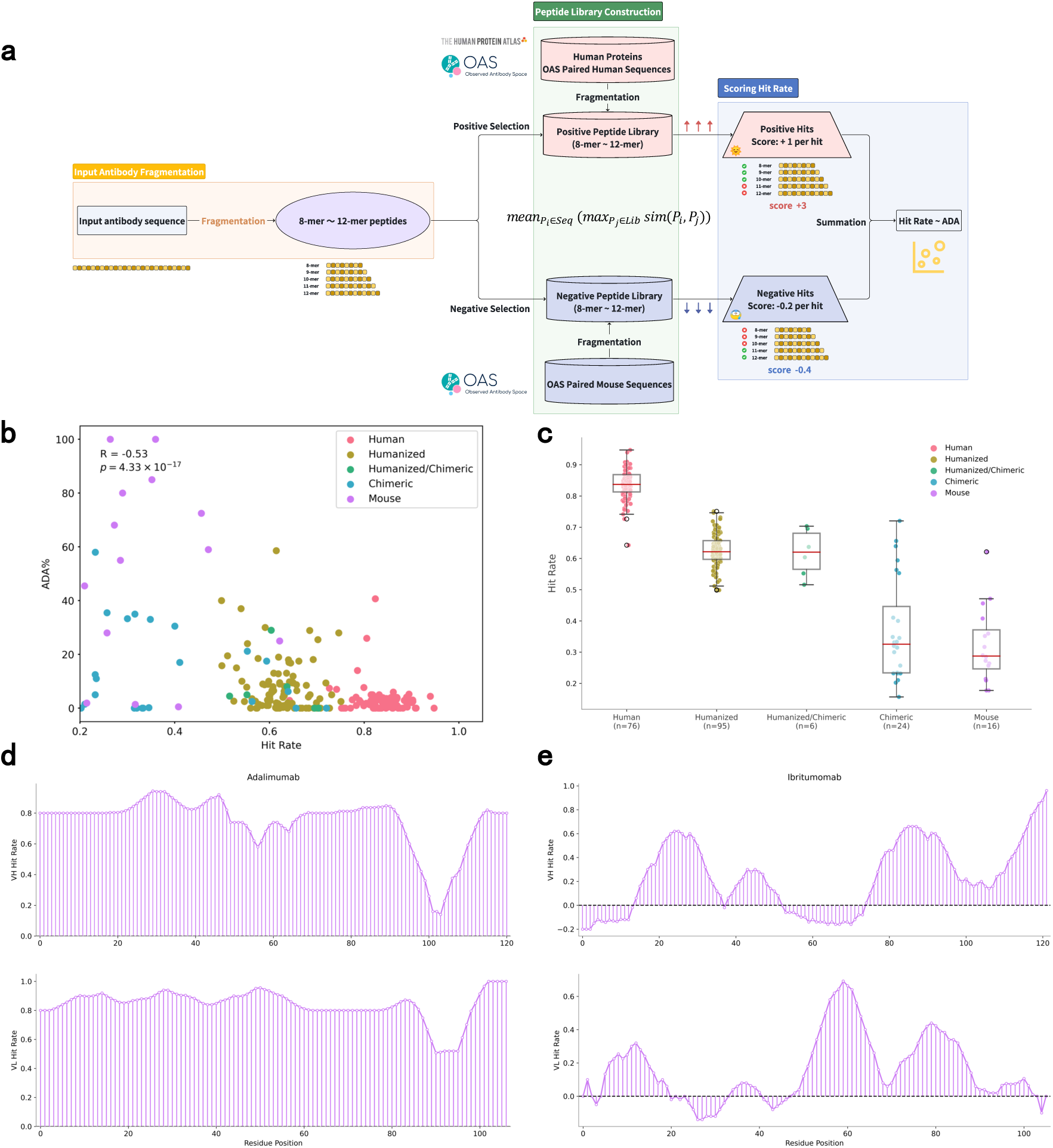
Overview of the ImmunoSeq model. **(a)** ImmunoSeq architecture. The workflow involves three core steps: (i) fragmentation of input antibody sequences into 8-mer to 12-mer peptide fragments (matching MHC-presented peptide lengths); (ii) dual-library selection, where fragments are compared against a positive peptide library (derived from human proteins and OAS human antibody sequences, 8-mers to 12-mers) and a negative peptide library (derived from OAS mouse antibody sequences, 8-mers to 12-mers); and (iii) hit rate scoring, where positive hits (matches to the positive library) contribute +1 per hit, negative hits (matches to the negative library) contribute -0.2 per hit, and summation yields the final hit rate, which correlates with clinical ADA risk. **(b)** Correlation between hit rate and clinical ADA risk. Scatter plot of hit rate vs. ADA incidence (%) for 217 therapeutic antibodies. Each point represents one antibody, colored by species origin. Pearson correlation coefficient (*R* = -0.53) and two-sided *P* value (*p* = 4.33×10⁻¹⁷) are shown (top left), indicating a significant negative correlation (higher hit rate indicates lower ADA risk). **(c)** Hit rate distribution across antibody species. Box plot of hit rate for 217 therapeutic antibodies from diverse species, colored by species origin which is consistent with panel **(b)**. Boxes represent interquartile range (IQR), horizontal lines denote medians, whiskers extend to 1.5×IQR, and individual points are outliers. **(d, e)** Residue-specific hit rate profiles. Residue-level hit rate distributions for **(d)** human antibody adalimumab (low immunogenicity) and **(e)** mouse antibody ibritumomab (high immunogenicity). Heavy chain variable region (VH) is shown in the top panel; light chain variable region (VL) is shown in the bottom panel. X-axis: residue index; Y-axis: residue-specific hit rate.

### Clinical ADA Validation

To quantify the performance of ImmunoSeq, we first evaluated the correlation between clinical ADA incidence and hit rate using 217 therapeutic antibodies. ImmunoSeq exhibits a robust negative correlation between hit rate and ADA incidence across 217 therapeutic antibodies, with a Pearson correlation coefficient (*R*) of -0.53 (*P* ≅ 4 × 10^!“#^) and a Spearman correlation coefficient (*ρ*) of - 0.41, demonstrating high consistency with immunogenicity (Fig. 1b). Antibodies scoring >0.8 exhibit median ADA rates below 5%, whereas those scoring <0.4 show median ADA incidence exceeding 40%, establishing semi-quantitative ADA incidence thresholds (Fig. 1c). This correlation is more obvious when focusing on human and mouse antibodies (Supplementary Fig. 1a). The median hit rate of human antibodies is 0.83, while the median hit rate of mouse antibodies is only 0.34, which is very different (Supplementary Fig. 1b).

This robust correlation further enables precise humanness classification on 553 therapeutic antibodies (Fig. 2b), in which ImmunoSeq can correctly discriminate between 198 human antibodies and 355 antibodies from other species (humanized, chimeric, mouse, felinized and caninized). Human antibodies achieve the highest hit rate, again aligning with minimal observed ADA induction. Humanized antibodies occupy an intermediate risk tier, corresponding to reduced immunogenicity after complementarity-determining regions (CDR) grafting. Chimeric and mouse antibodies display lowest hit rate, concordant with high clinical ADA rates (Fig. 2c and Supplementary Fig. 1c).

**Fig. 2.**
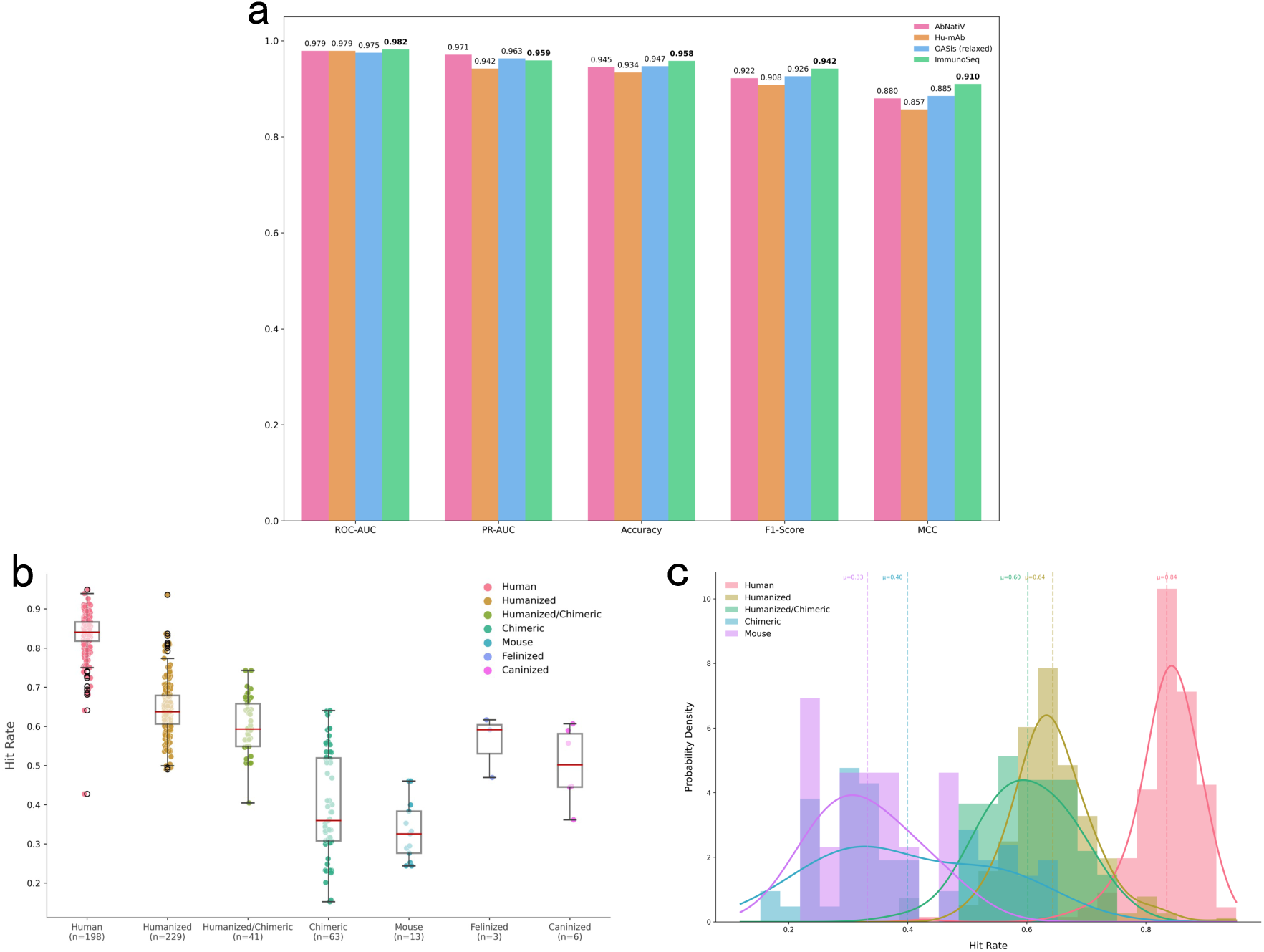
ImmunoSeq performance in humanness classification. **(a)** Benchmark comparison of classification metrics. ImmunoSeq was benchmarked against other models on a dataset of 553 therapeutic antibodies, tasked with distinguishing 198 human-derived therapeutics from 355 non-human-derived therapeutics (mouse, chimeric, and humanized). Metrics reported include ROC-AUC, PR-AUC, accuracy, F1-Score, and MCC, with ImmunoSeq outperforming comparative models across all metrics except PR-AUC. **(b)** Box plot of hit rate distribution by species. Hit rate distributions of antibodies from different species are shown, with boxes representing IQR, horizontal lines denoting medians, whiskers extending to 1.5×IQR, and points indicating outliers. **(c)** Bar plot of hit rate population by species. Bar heights represent the relative population of antibodies within each species category, with hit rate on the x-axis and species-colored bars reflecting the distribution of hit rates across species.

In benchmarking against established tools (AbNatiV, Hu-mAb, OASis), ImmunoSeq shows the best performance on ADA prediction (Table 1), with the highest *R* and *ρ* except *R* of Hu-mAb (*R*=-0.58). However, Hu-mAb is a random forest classifier trained supervisely to distinguish between human and mouse sequences, and its score is neither granular nor interpretable, which brings critical limitations. ImmunoSeq also demonstrated superior predictive capacity on humanness classification (Table 1 and Fig. 2a), with the area under the receiver operating characteristic curve (ROC-AUC) of 0.982 and Matthews correlation coefficient (MCC) of 0.910, surpassing all other methods. It should be noted that the area under the precision-recall curve (PR-AUC) of ImmunoSeq (PR-AUC=0.959) is slightly lower than that of AbNatiV (PR-AUC=0.971) and OASis (PR-AUC=0.953). To further quantify the performance of ImmunoSeq, we additionally assessed the humanness classification ability using AbNatiV test datasets, including Human, Human Diverse > 5%, Rhesus, PSSM-generated and Mouse dataset, each of which comprises of 10,000 antibody sequences from heavy, lambda and kappa databases of the corresponding species. The heavy, lambda and kappa classification performance of ImmunoSeq is comparable to those of deep-learning models for Human versus PSSM-generated or Mouse (Supplementary Fig. 12 and Supplementary Table 1-6). However, for Human versus Rhesus classification task of heavy, lambda and kappa, all models perform poorly except for AbNatiV which performs relatively well in the heavy chain classification task, possibly due to highly homologous of rhesus to humans.

**Table 1.** Evaluation of the humanness classification and immunogenicity correlation tasks.

| Method | Humanness Classification |  |  |  |  | Immunogenicity |  |
| --- | --- | --- | --- | --- | --- | --- | --- |
|  | PR-AUC | ROC-AUC | Accuracy | MCC | F1-Score | Pearson | Spearman |
| AbNatiV | **0.971** | 0.979 | 0.945 | 0.880 | 0.922 | -0.488 | -0.356 |
| Hu-mAb | 0.942 | 0.979 | 0.934 | 0.857 | 0.908 | **-0.584** | -0.405 |
| OASis (relaxed) | 0.963 | 0.975 | 0.947 | 0.885 | 0.926 | -0.511 | -0.387 |
| ImmunoSeq | 0.959 | **0.982** | **0.958** | **0.910** | **0.942** | -0.530 | **-0.409** |

We then conducted systematic ablation study to explore the influence of library selection, which confirms the dual-library architecture’s biological validity. Increasing the immunogenic penalty for each non-self-peptide from -0.2 to -1.0 can greatly lower the overall performance. Removing non-self-peptide library diminishes ADA risk discrimination (ROC-AUC decreases from 0.982 to 0.974) by impairing identification of cross-reactive T-cell epitopes (Extended Data Table 1). Further removal of human paired antibody sequences from self-peptide library significantly degrades both ADA prediction and classification performance (ROC-AUC collapses from 0.974 to 0.923; *ρ* collapses from -0.409 to -0.351), establishing that affinity-matured human antibodies incorporate critical tolerance signatures absent in the broader proteome. Notably, removal of human proteome sequences from self-peptide library causes a little influence on the performance, probably due to the relative smaller number of human proteome sequence. We also decomposed the hit rate for each peptide length, which suggested that different peptide length contribute and complement each other (Extended Data Table 2). Collectively, these results validate that 1) human antibody-derived peptides encode non-redundant tolerance determinants, 2) murine motifs provide essential negative controls for immunogenic hotspot identification, and 3) penalty calibration is required to balance library size effects without compromising biological fidelity.

### Site-specific Hit Rate Population

By calculating the average hit rate across all peptide fragments containing each residue, ImmunoSeq enables residue-specific immunogenicity assessment and pinpoints immunogenic hotspot residues. Analysis of residue-level profiles across 217 therapeutic antibodies reveals that human antibodies exhibit universally high hit rates: for example, most residues in both the heavy and light chains of adalimumab have hit rates above 0.8, whereas mouse antibodies show predominantly low hit rates (Fig. 1d). Ibritumomab shows frequent negative hit rates, particularly in CDRs, indicating residual murine motifs after CDR grafting onto human framework regions (FRs) - a pattern predictive of high immunogenicity risk (Fig. 1e). Further analysis of 25 humanization cases, where both precursor and humanized sequences were available and mutations localized to FRs, demonstrated substantial hit rate improvements upon humanization, with an average increase of 0.3, and humanized sequences exceeded hit rates of 0.7 (Fig. 3a). Residue-level analysis attributes these gains primarily to FRs (Supplementary Fig. 3-8), though adjacent CDRs also benefited, as exemplified by refanezumab, which showed hit rate improvements concentrated in FRs of both chains, with slight gains in CDRs (Fig. 3b,c). Assessment of ImmunoSeq’s application to nanobody humanization included eight nanobodies that neutralize the severe acute respiratory syndrome coronavirus 2 (SARS-CoV-2) spike protein receptor binding domain (RBD)^47^. Humanized mutations previously recommended by Llamanade involved 7 FR mutations^48^. Affinities remained unchanged after humanization, while ImmunoSeq predicted marginal hit rate improvements (∼0.1) confined to FRs, with CDRs retaining their original profiles - consistent with Llamanade’s findings (Fig. 3d,e). In conclusion, ImmunoSeq overcomes critical limitations of existing methods through residue-level hit rate mapping that localizes immunogenic hotspots for rational engineering, without requiring MHC haplotyping data, distinguishing it from “black-box” DL models via transparent, mechanism-driven scoring that directly reflects immune tolerance biology. This positions ImmunoSeq as a versatile tool that not only predicts immunogenicity but also enables precision engineering of antibody sequences.

**Fig. 3.**
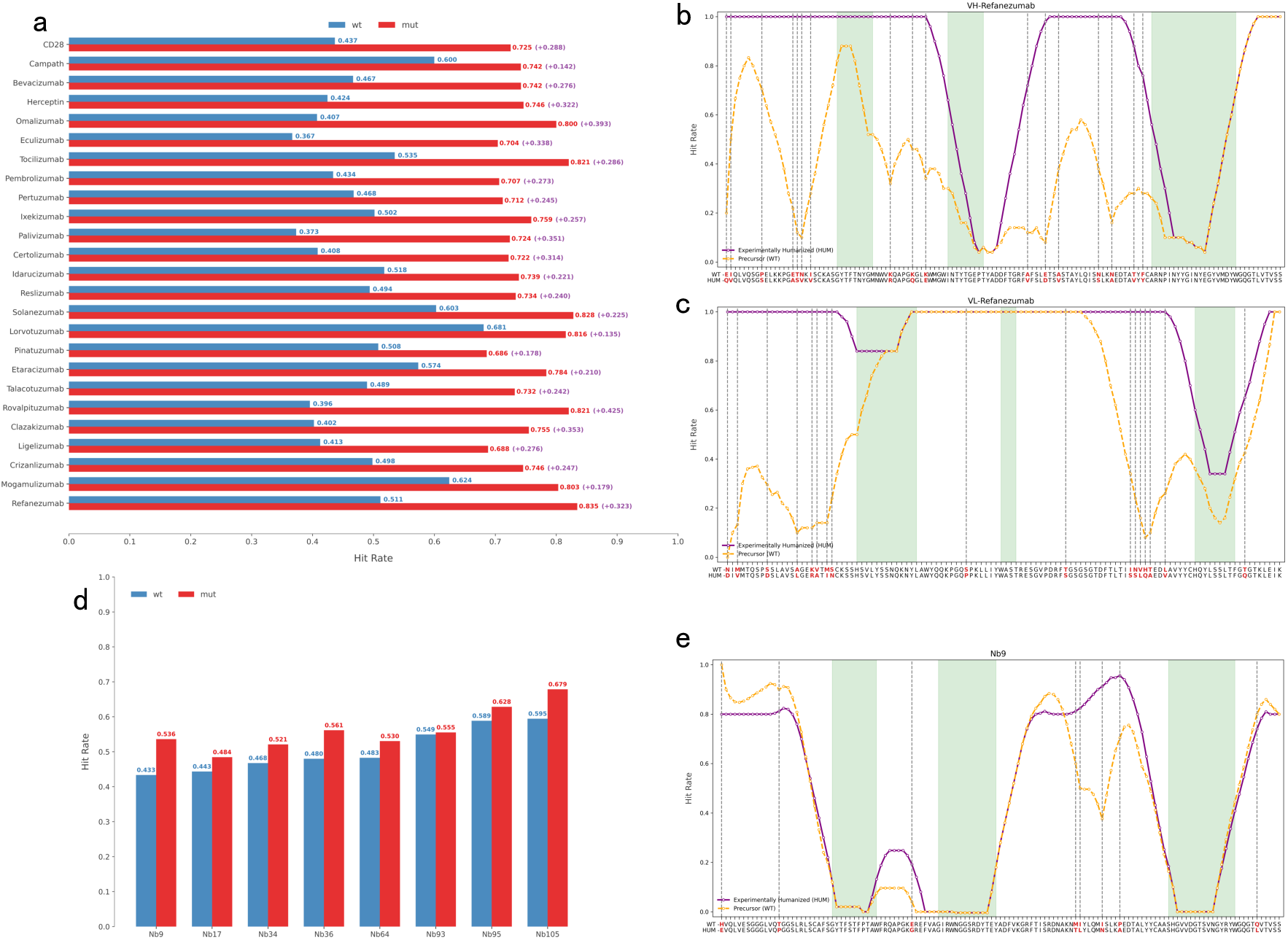
Prediction of ADA risk reduction upon humanization using ImmunoSeq. **(a)** Hit rate improvement in 25 humanized antibody pairs. Scatter plot comparing hit rates of wild-type (WT, blue) and humanized (red) sequences for 25 antibodies. All humanized sequences show increased hit rates (average improvement +0.3, range +0.135 to +0.425), with individual improvements labeled (e.g., Refanezumab: +0.323; Solanezumab: +0.225). **(b, c)** Site-specific hit rate profiles of Refanezumab. Residue-level hit rates for **(b)** heavy chain (VH) and **(c)** light chain (VL) of Refanezumab. CDR regions are shaded green; mutation sites are marked red. Yellow line: WT hit rate; purple line: humanized hit rate. Humanization mutations preferentially increase hit rates in framework regions (FRs), with minimal impact on CDRs. **(d)** Hit rate improvement in 8 SARS-CoV-2 nanobodies. Scatter plot comparing WT (blue) and humanized (red) hit rates for 8 nanobodies (Nb9, Nb17, Nb34, Nb36, Nb64, Nb93, Nb95, Nb105). All humanized nanobodies show increased hit rates, consistent with reduced ADA risk. **(e)** Site-specific hit rate profile of Nb9. Residue-level hit rate distribution for nanobody Nb9 (WT: yellow line; humanized: purple line), with CDR regions shaded green and humanization mutation sites marked in red. Mutations in FR2 and FR3 drive hit rate improvement (+0.103).

### Sequence Humanization Design

We performed *in silico* experiments using the above 25 humanization cases to evaluate the ability of ImmunoSeq to recall experimentally validated humanization mutations from precursor sequences, a task that serves as a standard for immunogenicity optimization. Notably, experimental mutations may not represent the only ground truth, and certain substitutions may fail to reduce or even paradoxically elevate immunogenicity risk through emergent epitope presentation, structural perturbation, or colloidal instability; nevertheless, the observed consistency between ImmunoSeq recommendations and experimentally validated mutations substantiates its reliability. We leveraged two distinct recommendation strategies: (i) one-shot design, where all possible mutations are ranked by their hit rates, and (ii) iterative design, where the top-ranked mutation from previous rounds is iteratively incorporated into parent sequences and re-evaluated in subsequent rounds - providing interpretable mutation evolutionary trajectories. For the first strategy, we quantified two metrics: (1) the proportion of experimental mutations that increase hit rate relative to precursor sequences, and (2) the top-K (K=1,5,10,20,40) enrichment ratio of experimental mutations in recommendations.

We found that more than half of the experimental mutations increased hit rate in 21 out of 25 antibodies (Fig. 4a), with 7 out of 25 antibodies exhibiting >80% hit rate-increasing mutations (e.g., Solanezumab, where all 26 experimental mutations increased hit rate). We also observed mutations that reduced hit rates, likely corresponding to the above immunologically ineffective or unexpectly risk-elevating substitutions. Further evaluation of top-K enrichment ratios yielded a top-1 ratio of 0.60, top-5 of 0.51, top-10 of 0.48, top-20 of 0.38, and top-40 of 0.33 (Fig. 4b), with the top 1 or 2 mutations for most antibodies being experimentally validated and exhibiting the highest hit rate (Fig. 4c). To explore structural rationales, we analyzed three cases (Solanezumab, Tocilizumab, Pembrolizumab) by focusing on experimental mutations within the top-10 recommendations (Fig. 4d-f). The locations of these mutations aligned with established principles - where FR mutations prioritize structural stability and CDR conformation maintenance - with top-ranked mutations enriched in surface-exposed FR residues and avoiding core conserved residues, indicating the model implicitly balances “surface immunogenicity reduction” and “structural stability preservation,” mirroring empirical design rules. For the second strategy, which targets a more practical scenario, iterative design showed hit rate increasing monotonically with rounds, converging by around 30 rounds to a mean hit rate of 0.84 (Fig. 5a), exceeding the mean hit rate of experimental humanized sequence (0.75). It also improved top-K enrichment ratios (top-1: 0.60; top-5: 0.65; top-10: 0.61; top-20: 0.57; top-40: 0.49), outperforming one-shot design with larger gains in later rounds (Fig. 5b and Supplementary Fig. 9-11). Four antibodies (Ixekizumab, Ligelizumab, Mogamulizumab, and Talacotuzumab) achieved >0.6 enrichment rates after 40 rounds (Fig. 5c-f), with hit rate surpassing experimental sequences, while a minority (Crizanlizumab, Herceptin, Pinatuzumab, Refanezumab, Solanezumab) showed lower hit numbers in iterative design than one-shot design, likely due to convergence to local optimal. To further benchmark the efficiency of humanization design, we collected optimized sequences from Hu-mAb and Sapiens and calculated their hit rates (Supplementary Table 7). The average hit rate of ImmunoSeq consistently outperforms those of other methods, while the number of recommended mutations are comparable to those suggested by other models. Collectively, ImmunoSeq’s top-ranked mutations strongly correlated with experimental humanization mutations, aligned with structural design principles, and iterative optimization further enhanced performance, supporting its utility in rational antibody engineering.

**Fig. 4.**
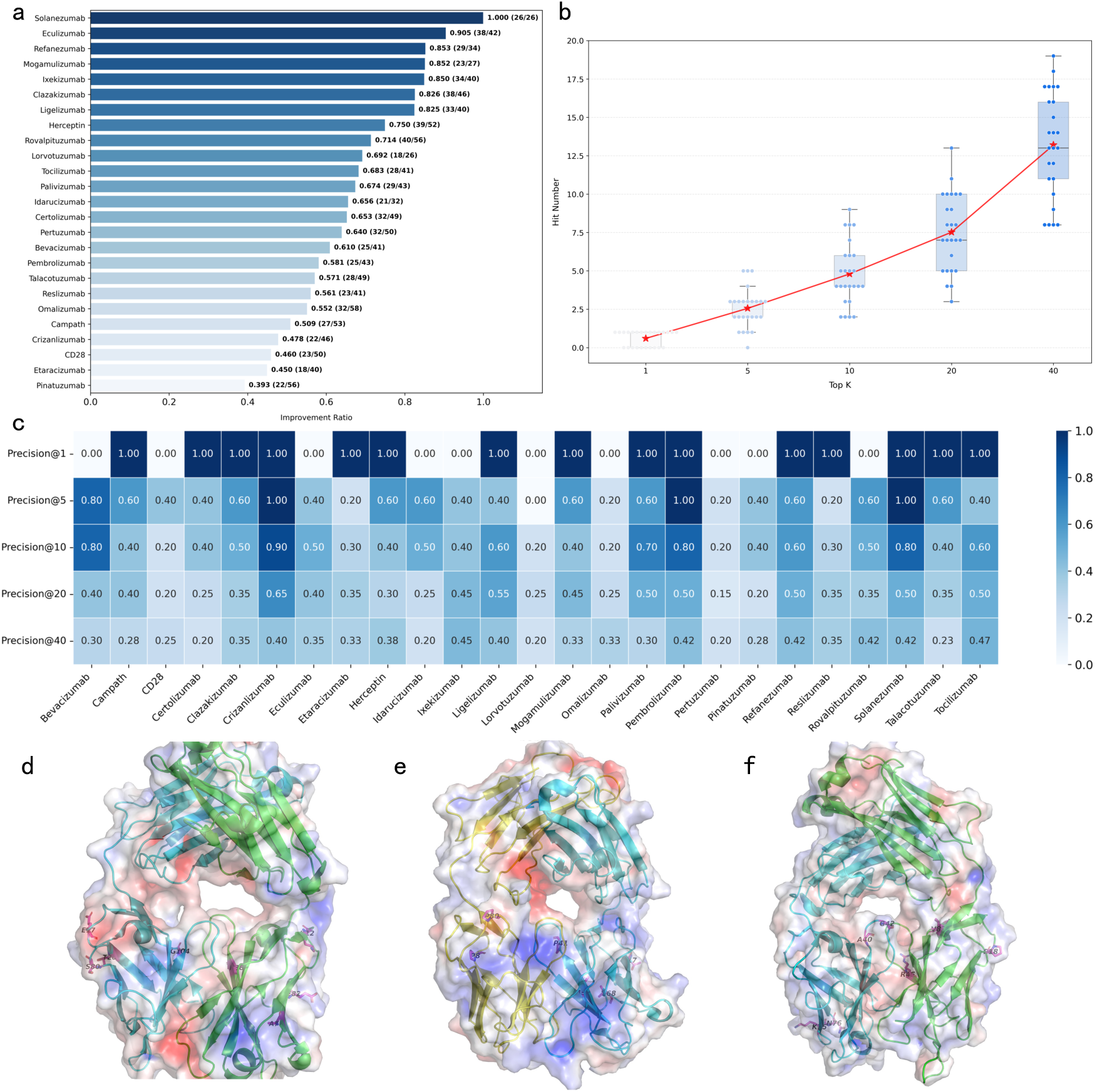
Humanization design using the one-shot strategy. **(a)** Hit rate improvement ratio of experimentally validated humanization mutations. Bar plot showing the proportion of humanization mutations that increase hit rate for 25 antibodies. Each bar corresponds to one therapeutic antibody, labeled with its name and improvement ratio (mutations increasing hit rate/total mutations). Over half of the mutations increased hit rate in 21/25 antibodies (median improvement ratio: 0.653). **(b)** Top-K precision of experimental mutation enrichment. Line plot showing precision vs. top-K recommendations (K=1,5,10,20,40). Precision values are: Precision@1=0.60, Precision@5=0.51, Precision@10=0.48, Precision@20=0.38, Precision@40=0.33, indicating strong enrichment of experimentally validated mutations in top-ranked recommendations. **(c)** Heatmap of precision@K across 25 antibody cases. Precision values for top-1, top-5, top-10, top-20, and top-40 mutations are shown for each antibody (rows). 15 out of 25 antibodies exhibit precision=1.00 for top-1 mutations, with precision decreasing for larger K. **(d-f)** Structural analysis of recommended mutations for **(d)** Solanezumab, **(e)** Tocilizumab, and **(f)** Pembrolizumab. Top-10 overlapping mutations (computational prediction vs. experimental validation) are shown as sticks. Proteins are displayed as cartoons with electrostatic surfaces (blue: positive charge; red: negative charge). Mutations are enriched in surface-exposed FR residues and avoid core conserved residues, aligning with structural stability principles. The overlapping residues are listed below: Solanezumab VH: A40, G42, K75, N76; VL: P18, R45, A80, V83. Tocilizumab VH: P8, P80; VL: T17, P41, I49, V68. Pembrolizumab VH: K12, R38, A79, E82; VL: E17, T20, S80, G104.

**Fig. 5.**
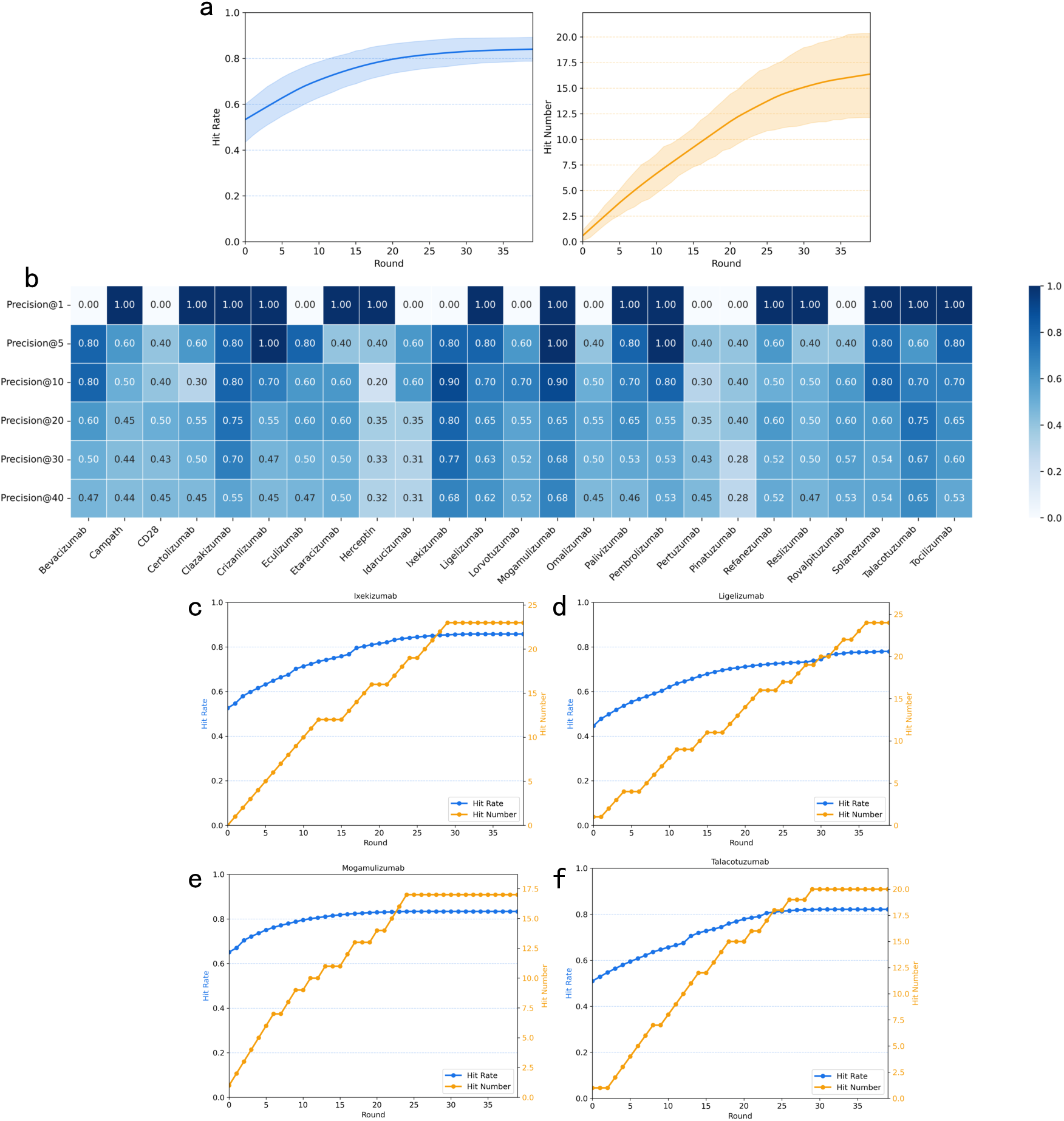
Humanization design using the greedy iterative strategy. **(a)** Convergence of hit rate and hit number during iterative optimization. Line plots showing hit rate (blue) and hit number (orange) vs. iteration rounds (0-39). Both metrics gradually increase and converge at ∼30 rounds, reaching a median hit rate of 0.84 and median hit number of 17.5. These values are significantly higher than the average hit rate of experimental humanized sequences (0.75) and the hit number of the one-shot strategy (12.5). **(b)** Top-K enrichment ratio across 25 humanization cases. Heatmap of precision@K (K=1,5,10,20,40) for the greedy strategy, with each row representing one antibody. Median precision values are as follows: precision@1=0.60, precision@5=0.65, precision@10=0.61, precision@20=0.57, precision@40=0.49. These values are consistently higher than those of the one-shot strategy (Fig. 4b). **(c-f)** Detailed iterative optimization of three antibodies. **(c)** Ixekizumab, **(d)** Ligelizumab, **(e)** Mogamulizumab, and **(f)** Talacotuzumab. Hit rate (blue) and hit number (orange) curves demonstrate steady improvement over iterations, converging at 25-30 rounds. For Ixekizumab: hit rate=0.858 (vs. experimental humanized hit rate 0.759) and hit number=23 (vs. one-shot hit number 18). For Ligelizumab: hit rate=0.780 (vs. experimental humanized hit rate 0.688) and hit number=24 (vs. one-shot hit number 16). For Mogamulizumab: hit rate=0.834 (vs. experimental humanized hit rate 0.803) and hit number=17 (vs. one-shot hit number 13). For Talacotuzumab: hit rate=0.821 (vs. experimental humanized hit rate 0.732) and hit number=20 (vs. one-shot hit number 9). All metrics exceed those of both experimental humanized sequences and the one-shot strategy.

## Discussion

In this work, we introduce ImmunoSeq – an immunogenicity prediction model that translates human antibody evolutionary principles into a computational framework via dual-library peptide matching. ImmunoSeq achieves robust ADA correlation (R=-0.53; *ρ*=-0.41) across a dataset of 217 clinical antibodies and superior humanness classification performance on 553 therapeutic antibodies, outperforming DL models. Ablation studies confirm the critical importance of the dual-library architecture. Residue-level hit rate profiling enables precise mapping of site-specific immunogenicity contributions, successfully predicting ADA changes upon humanization while identifying immunogenic hotspots to guide engineering. Complementing these capabilities, the design protocol achieves superior enrichment of experimental mutations (top-1: 0.6) through interpretable one-shot design and traceable iterative optimization that converges to a hit rate of 0.84, exceeding empirically humanized sequences.

To enhance ImmunoSeq’s immunogenicity optimization capabilities and bridge computational predictions to biophysical scenario, structural information can be integrated into the framework. The thermostability and aggregation propensity of optimized sequences can further be evaluated using Rosetta^49–50^, AlphaFold3^51^, and molecular dynamics simulations^52^ to filter out structurally unstable and aggregation-prone variants. Critically, binding affinity preservation during humanization can be validated through free energy calculation methods^53–58^ including free energy perturbation and Rosetta cartesian_ddG, establishing a multi-parameter design funnel prioritizing hit rate → structural integrity → binding affinity. This workflow positions ImmunoSeq as the computational engine of an integrated antibody engineering pipeline that feeds optimized sequences into downstream experimental validation.

Despite these successes, three key limitations merit consideration: First, the sequence-only approach ignores structural epitopes despite substantial differences in surface features between human and murine antibodies. Integrating surface similarity metrics (e.g., using MaSIF^59–61^) would project residue-level structural information into topological space to better measure human/murine similarity, potentially improving ADA correlations and humanness classification. This could be implemented through a structural module that predicts 3D structures, extracts topological surface features, and scores human/murine similarity. Second, the scoring system requires refinement: a)

Introducing protease-specific cleavage probability weighting for k-mers via protease-site prediction, as not all sites are cleaved by proteases; b) Broadening hit-matching criteria beyond sequence identity between input and library peptides (e.g., edit distance, protein language model embeddings); c) Implementing threshold-based risk focusing that excludes high hit-rate (>0.7) peptides, since ADA risk primarily stems from low hit-rate regions. These aspects present opportunities to integrate DL modules for cleavage site prediction, surface feature extraction, and refined scoring functions. Third, the efficiency of iterative design is hampered by greedy optimization that may converge to local optima. Future implementations could incorporate Monte Carlo Tree Search to explore global mutational space and identify optimal combinatorial solutions.

## Methods

### Datasets and Peptide Library Construction

Human proteomic sequences were retrieved from the AlphaFold Protein Structure Database, comprising 23,391 entries. After removing duplicates, 21,931 unique human protein sequences were retained. Human antibody sequences were sourced from the Observed Antibody Space (OAS) Paired Database, which was filtered to include only sequences from healthy donors to exclude exogenous sequences associated with potential ADA risk, yielding 1,177,822 sequences. Murine antibody sequences were similarly retrieved from the OAS Paired Database, resulting in 84,449 sequences. Filtered human protein and antibody sequences were used to construct the positive peptide library, whereas filtered murine antibody sequences formed the negative peptide library. For each sequence, all overlapping k-mer peptides (k=8-12) were extracted, and duplicate peptides were removed. This process generated >700 million positive peptides and 40 million negative peptides. To efficiently store and query these large peptide datasets, bloom filters were implemented, substantially reducing memory usage and accelerating peptide match queries.

### ImmunoSeq Scoring Method

Generally, the hit rate of a query antibody sequence against a peptide library is formally defined as:

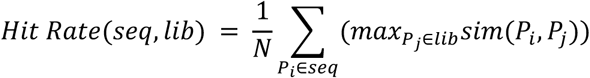

where *seq* denotes the query antibody sequence, *lib* denotes the peptide library, and *N* is the total number of k-mer fragments extracted from *seq*. The similarity function *sim*(*P_i_*, *P_j_*) was defined as an identity metric: *sim*(*P_i_*, *P_j_*) = 1 if *P_i_* and *P_j_* are identical k-mer sequences, and 0 otherwise. The sum of similarity scores was normalized by *N* to yield the average hit rate for the query sequence against the library.

To account for differences in library size and balance contributions from self vs. non-self-peptides, the final hit rate of the query sequence was computed as:

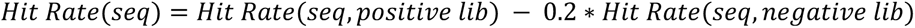

Here, *positive lib* and *negative lib* refer to the positive self-peptide library and negative non-self-peptide library, respectively. The scaling factor of 0.2 for the negative library was empirically determined to balance the larger size of the positive library relative to the negative library. For antibody sequences, hit rates were computed separately for the heavy chain and light chain, then averaged to generate a global hit rate representing the overall self-similarity of the antibody. This global hit rate was used to quantify correlations with ADA incidence and classify humanness.

To enable residue-level immunogenicity profiling, we further computed local hit rates for individual residues. For each residue in the antibody sequence, all overlapping k-mer fragments containing that residue were extracted, and their hit rate against the positive/negative libraries was calculated using the above formula. This residue-specific hit rate population identifies immunogenic hotspots (residues with low local hit rates) to guide sequence optimization.

### Performance Metrics

To assess the relationship between ImmunoSeq derived hit rates and clinical ADA risk, we computed Pearson correlation coefficient (*R*) and Spearman correlation coefficient (*ρ*). Pearson’s *R* measures the linear association between two continuous variables, ranging from -1 (perfect negative linear correlation) to 1 (perfect positive linear correlation), which is defined as:

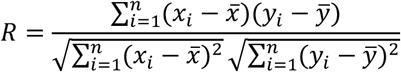

where *n* is the total number of samples, *x_i_* and *y_i_* are individual observations of the two variables, and *x̄*, *ȳ* G are their respective means. The numerator represents the covariance of *x* and *y*, and the denominator is the product of their standard deviations.

Spearmans’s *ρ* assesses monotonic relationship between two variables by ranking observations. It is equivalent to Pearson’s *R* computed on the ranks of *x* and *y* (denoted as *R*_0_ and *R*_1_). Unlike Pearson’s *R*, Spearmans’s *ρ* is robust to outliers and non-linear trends.

For humanness classification tasks (e.g., distinguishing human vs. non-human antibodies), we quantified performance using ROC-AUC, PR-AUC, accuracy, F1-score, and Matthew’s correlation coefficient (MCC).

ROC-AUC quantifies the model’s ability to distinguish classes by plotting true positive rate (TPR) against false positive rate (FPR) across thresholds. A perfect classifier has ROC-AUC = 1, while random performance yields ROC-AUC = 0.5.

Complementary to ROC-AUC, PR-AUC focuses on precision (positive predictive value) vs. recall, which is ideal for imbalanced datasets, with PR-AUC = 1 for perfect classification and 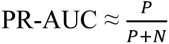 for random performance (*P*: positive samples, *N*: negative samples).

Accuracy is the proportion of correctly classified samples, defined as:

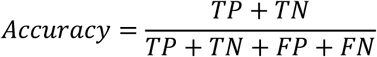

where *TP* (true positives) is correctly classified positive samples, *TN* (true negatives) is correctly classified negative samples, *FP* (false positives) is negative samples misclassified as positive, and

*FN* (false negatives) is positive samples misclassified as negative.

F1-Score balances precision and recall, which is ideal for imbalanced datasets and defined as:

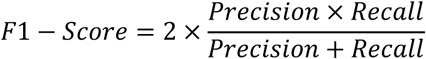

where precision equals to 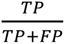 and recall equals to 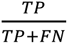.

MCC is a robust metric for binary classification, which incorporates all elements of the confusion matrix and ranges from -1 (perfect misclassification) to 1 (perfect prediction), with 0 indicating random performance. It is particularly valuable for imbalanced datasets, as it avoids bias toward majority classes, defined as:

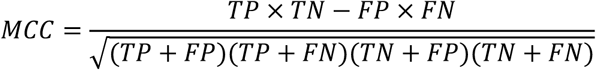

### Benchmarking with Other Methods

To validate ImmunoSeq’s performance, we conducted two benchmarking tasks: (1) ADA risk correlation using 217 therapeutic antibodies with clinical ADA incidence data; and (2) humanness classification using 553 therapeutic antibodies from the BioPhi dataset.

For humanness classification, metrics like F1-score, accuracy, and MCC depend on the hit rate threshold (cutoff for classifying sequences as human sequences). To enable fair comparison with other methods, we optimized this threshold for the BioPhi dataset: ImmunoSeq’s optimal hit rate threshold was determined to be 0.75 (sequences with hit rate above 0.75 were classified as human sequences; those below as non-human sequences). Notably, this threshold may vary with peptide library composition (e.g., size or sequence diversity of positive/negative libraries).

We compared ImmunoSeq to three state-of-the-art methods: AbNatiV, Hu-mAb, and OASis. AbNatiV is an antibody language model based on a vector quantized variational autoencoder (VQ-VAE) framework, trained unsupervised on human antibody sequences from the OAS database. Hu-mAb is a supervised random forest model trained to distinguish human vs. murine sequences using handcrafted sequence features. OASis is a sequence-based immunogenicity predictor that calculates 9-mer peptide similarity against the OAS database. Benchmark metrics for comparative methods were extracted from their original publications to ensure consistency with reported performance.

To further validate generalizability, we evaluated ImmunoSeq on the AbNatiV test dataset, which comprises 10,000 sequences per category: Human, Human Diverse (>5% sequence divergence from germline), Rhesus, PSSM-generated (synthetic non-human), and Murine. For comparative methods, we used metrics reported in the AbNatiV study to maintain consistency in evaluation protocol.

### Prediction on SARS-CoV-2 nanobody therapeutics

To further extend validation to nanobodies - single-domain antibodies with distinct framework region FR architecture - we evaluated ImmunoSeq’s ability to predict immunogenicity changes in 8 previously characterized SARS-CoV-2 neutralizing nanobodies. These nanobodies had undergone humanization via Llamanade, an open-source computational pipeline designed to guide nanobody humanization by introducing FR mutations. For each nanobody, Llamanade recommended 7 FR mutations targeting conserved residues, which were experimentally validated to preserve binding affinity to the SARS-CoV-2 spike protein receptor-binding domain (RBD). This humanized dataset provided a unique opportunity to test ImmunoSeq’s performance on non-canonical antibody formats (single domain vs. conventional IgG).

### Humanization Design Protocol

To evaluate ImmunoSeq’s ability to guide immunogenicity optimization, we used 25 antibody pairs (pre-humanization and post-humanization sequences) from the Hu-mAb dataset, each containing FR mutations introduced during humanization. Our design protocol aimed to maximize the overall hit rate of the input sequence by iteratively mutating FR residues (non-CDR positions), employing two distinct mutation sampling strategies: one-shot design and iterative design.

One-shot design strategy enumerates all possible single-point mutations across FR residues (each position mutated to the rest 19 natural amino acids), generating a library of mutated sequences. For each mutant, we computed the hit rate and ranked mutations by their ability to increase hit rate relative to the wild-type (WT) sequence. This approach efficiently identifies high-impact mutations in a single step, as it directly enriches for all FR mutations that boost hit rate without requiring iterative refinement.

In contrast, iterative design employs a greedy search algorithm with stepwise mutation incorporation:

1. Initialization: Start with the WT sequence and compute its baseline hit rate.
2. Mutation generation: Enumerate all single-point FR mutations and calculate their hit rate.
3. Selection: Select the top-ranked mutation (highest hit rate increase) and incorporate it into the sequence.
4. Convergence check: Repeat steps 2-3 with the updated sequence until hit rate plateaus.

This strategy traces the evolutionary trajectory of mutations, enabling residue-level interpretation of how each mutation contributes to hit rate improvement, thereby enhancing interpretability of the optimization process.

## Data Availability

All the data required to analyze the ADA incidence correlation, humanness classification, and immunogenicity optimization are provided in the Supplementary Data file.

## Code Availability

The ImmunoSeq code is available at https://github.com/qj-Huang/ImmunoSeq.

## Extended data

**Extended Data Table 1.** Ablation study on the influence of library selection.

| Sequence Library |  | Humanness Classification |  |  |  |  | Immunogenicity |  |
| --- | --- | --- | --- | --- | --- | --- | --- | --- |
| Self-peptide | Non-self-peptide | PR-AUC | ROC-AUC | Accuracy | MCC | F1-Score | Pearson | Spearman |
| Human |  | 0.901 | 0.923 | 0.879 | 0.734 | 0.813 | -0.510 | -0.351 |
| OAS Paired |  | 0.944 | 0.974 | 0.944 | 0.878 | 0.921 | -0.521 | -0.393 |
| Human + OAS Paired |  | 0.944 | 0.974 | 0.946 | 0.882 | 0.925 | -0.520 | -0.393 |
| OAS Paired | Mouse | 0.958 | 0.982 | 0.958 | **0.911** | **0.943** | -0.530 | -0.409 |
| **Human + OAS Paired** | **Mouse** | **0.959** | **0.982** | **0.958** | 0.910 | 0.942 | **-0.530** | **-0.409** |

**Extended Data Table 2.**
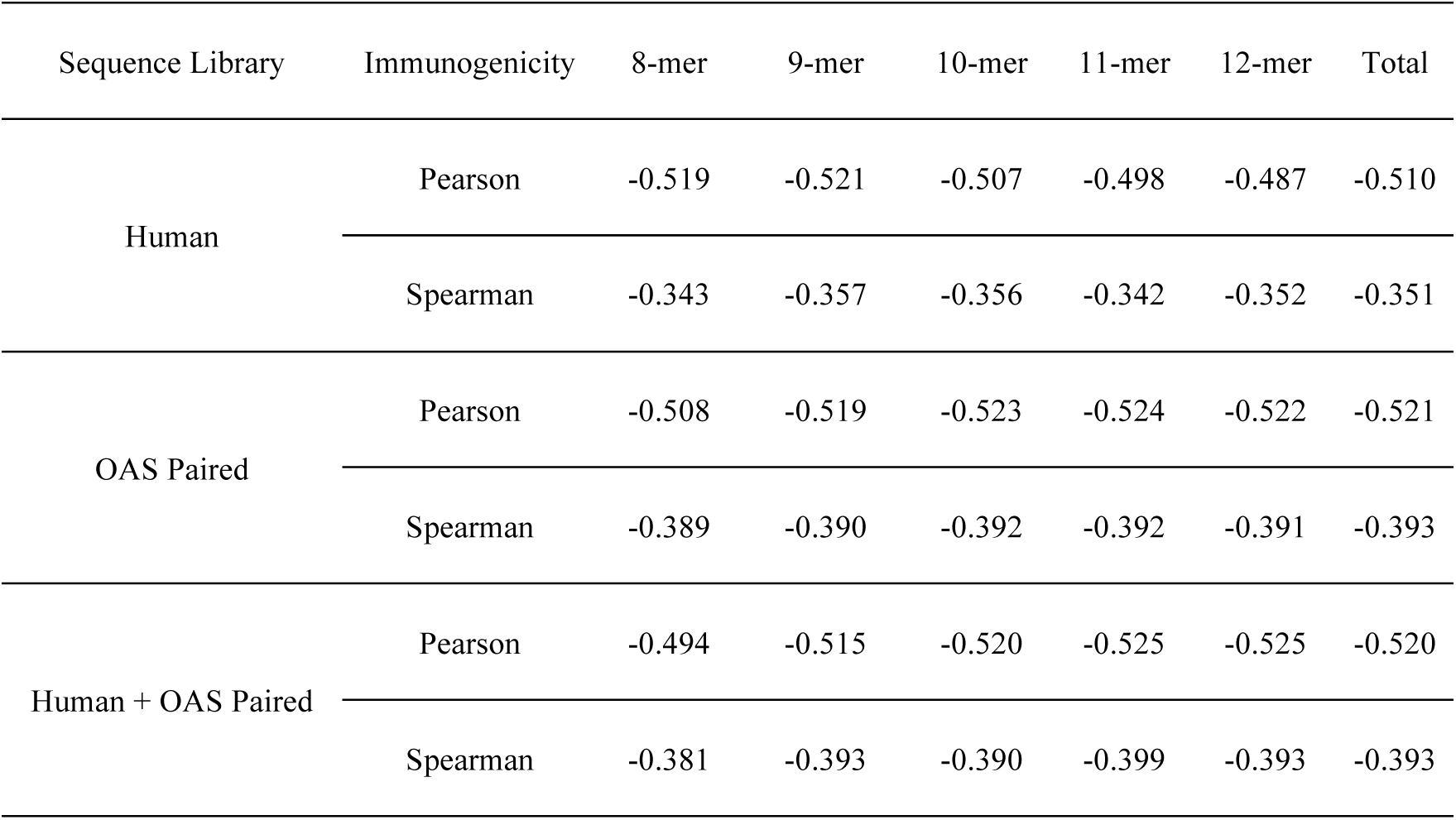
Hit rate decomposition for each k-mer peptides.

