## Supplementary Information for "Antibody immunogenicity prediction and optimization with ImmunoSeq"

### Table of Contents

**Supplementary Figures 1-12**

Supplementary Fig. 1. Performance of ImmunoSeq in classifying human and mouse antibodies.

Supplementary Fig. 2. Site-specific hit rate profiles of 8 SARS-CoV-2 nanobodies.

Supplementary Fig. 3 - Fig. 8. Site-specific hit rate profiles for heavy chain and light chain of therapeutics antibodies.

Supplementary Fig. 9 - Fig. 11. Humanization design using the greedy iterative strategy.

**Supplementary Tables 1-6**

Supplementary Table 1. Evaluation of the ROC-AUC classification task for human heavy chain sequences from AbNatiV

Supplementary Table 2. Evaluation of the ROC-AUC classification task for human lambda light chain sequences from AbNatiV

Supplementary Table 3. Evaluation of the ROC-AUC classification task for human kappa light chain sequences from AbNatiV

Supplementary Table 4. Evaluation of the PR-AUC classification task for human heavy chain sequences from AbNatiV

Supplementary Table 5. Evaluation of the PR-AUC classification task for human lambda light chain sequences from AbNatiV

Supplementary Table 6. Evaluation of the PR-AUC classification task for human kappa light chain sequences from AbNatiV

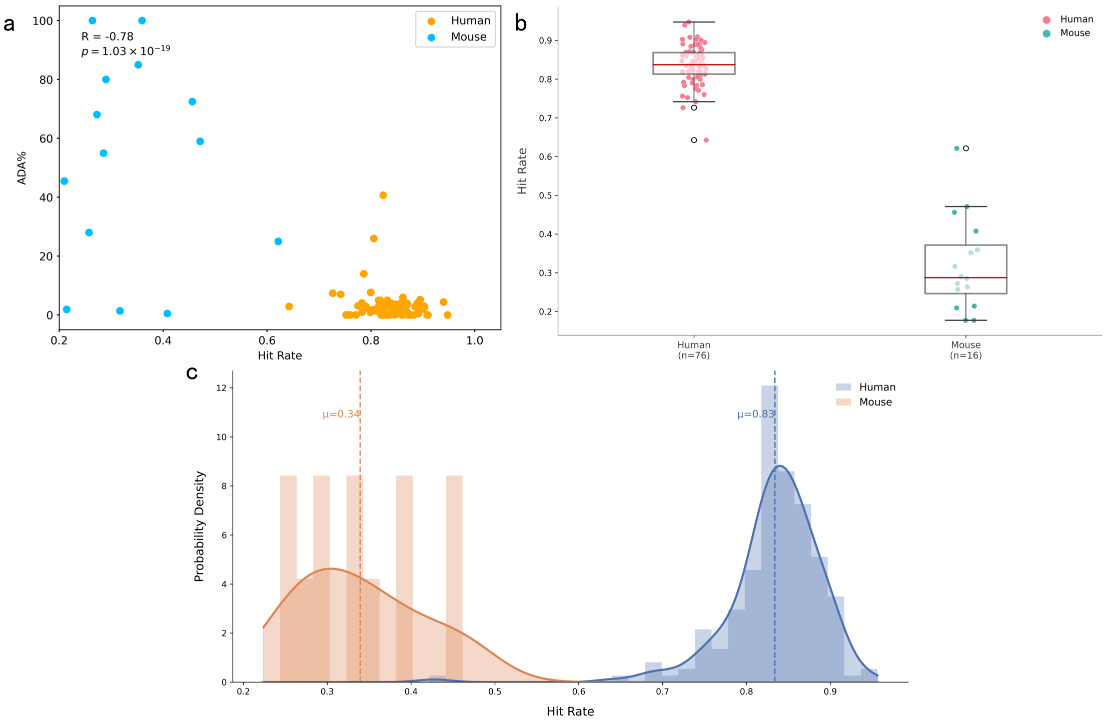


**Supplementary Fig. 1. Performance of ImmunoSeq in classifying human and mouse antibodies. (a)** Scatter plot of ADA incidence vs. hit rate in therapeutic antibodies. Scatter plot showing the correlation between ADA incidence and hit rate for human (yellow) and mouse (blue) antibodies from a dataset of 217 therapeutic antibodies. Each data point represents an individual therapeutic antibody. **(b)** Distribution of hit rates in human vs. mouse therapeutic antibodies. Box plot comparing the population distribution of hit rates between human and mouse antibodies, derived from the same 217 therapeutic antibody dataset as in **(a)**. Boxes represent interquartile ranges (IQRs), horizontal lines denote medians, whiskers extend to 1.5×IQR, and outliers are shown as individual points. **(c)** Distribution of hit rates for human and mouse antibodies from BioPhi dataset.

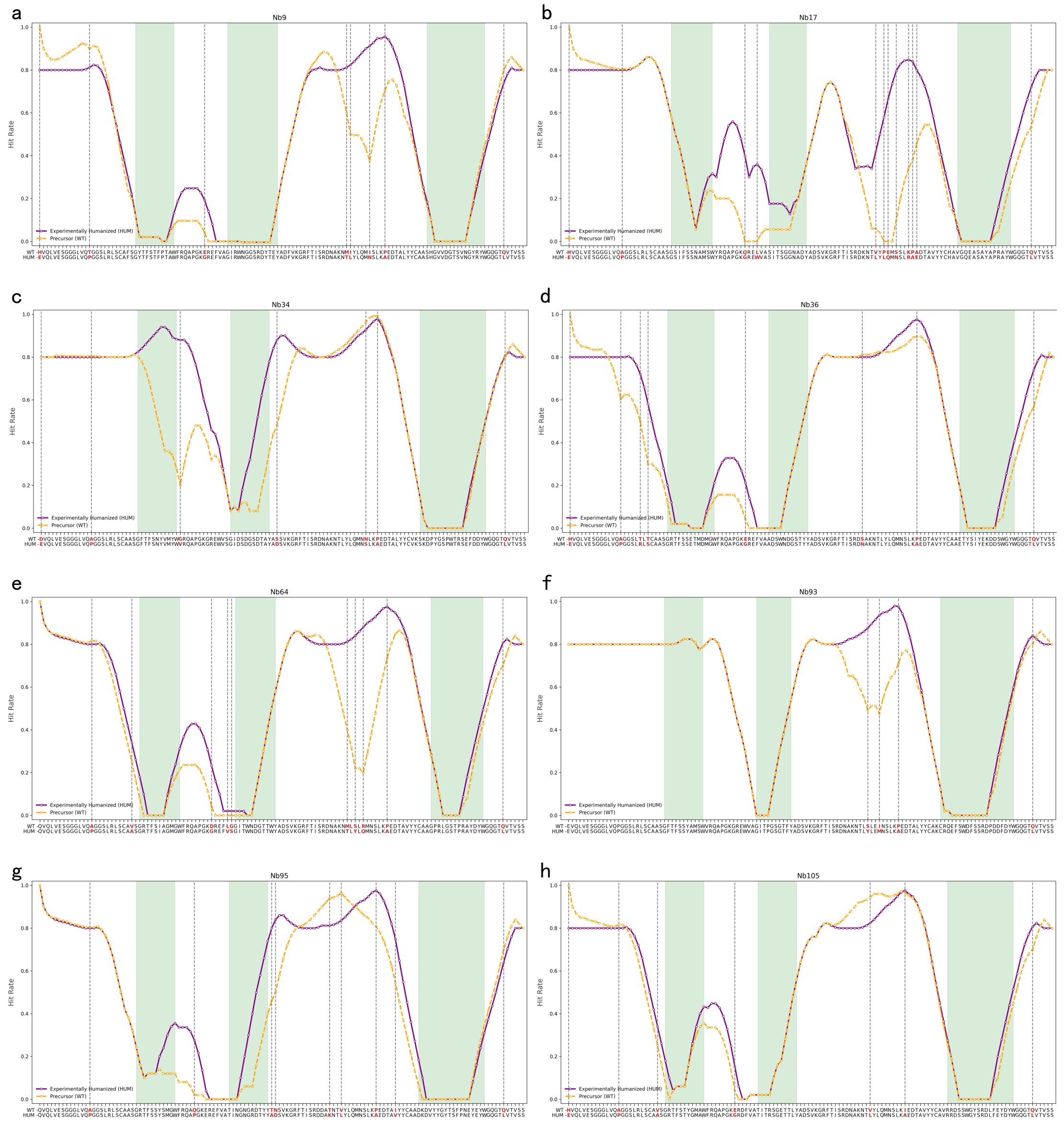


**Supplementary Fig. 2. Site-specific hit rate profiles of 8 SARS-CoV-2 nanobodies. (a)-(h)** correspond to nanobodies Nb9, Nb17, Nb34, Nb36, Nb64, Nb93, Nb95, and Nb105, respectively. Residue-level hit rate distribution for a representative nanobody Nb9 in panel **(a)**: wild-type (WT, yellow line) and humanized (purple line) variants. CDRs are shaded green; humanization mutation sites are marked in red.

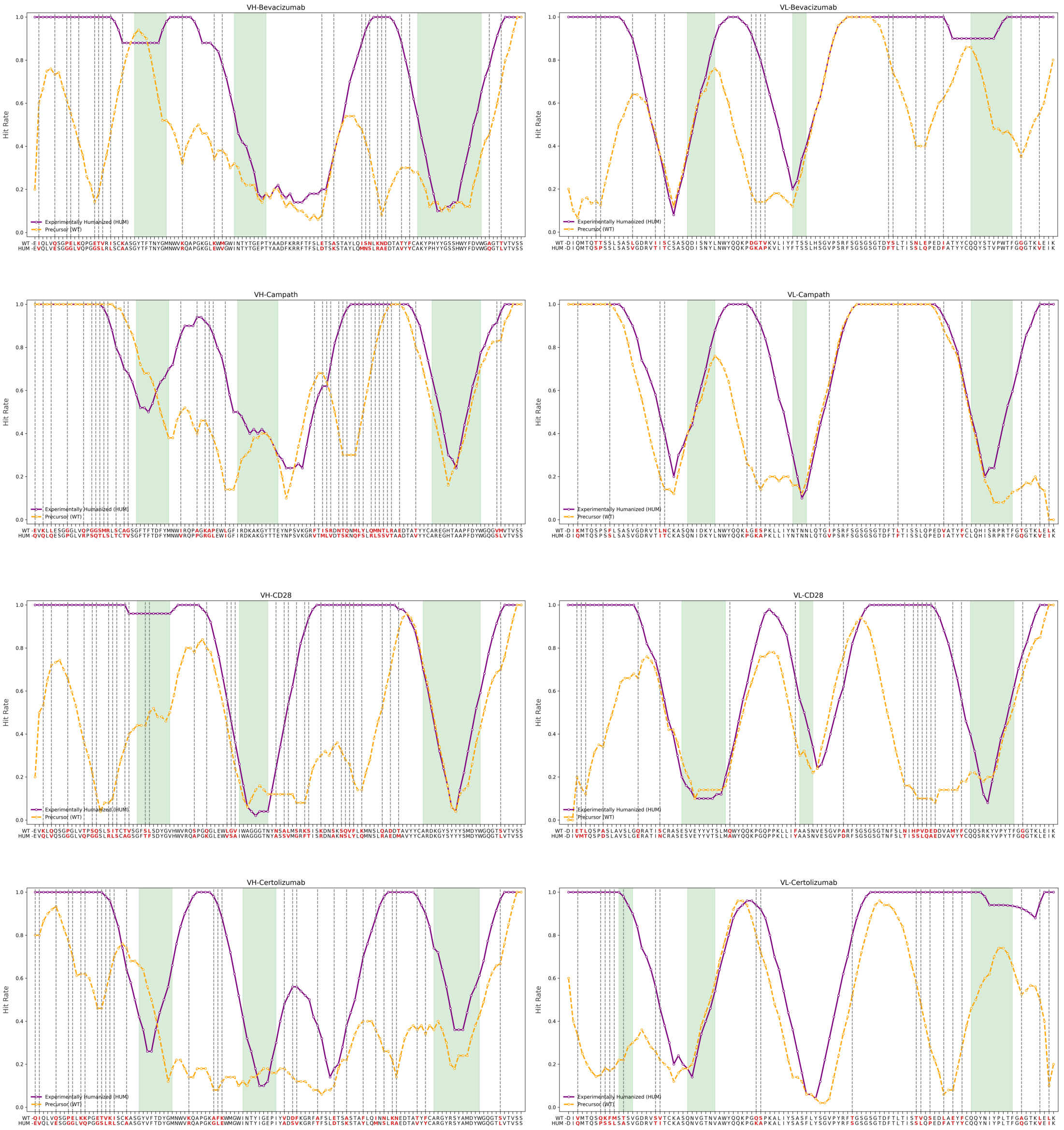

**Supplementary Fig. 3. Site-specific hit rate profiles for heavy chain (VH, left) and light chain (VL, right) of therapeutics antibodies.** From top to bottom: Bevacizumab, Campath, CD28, and Certolizumab. CDR regions are shaded green; mutation sites are marked red. Yellow line: WT hit rate; purple line: humanized hit rate. Humanization mutations preferentially increase hit rates in FRs, with minimal impact on CDRs.

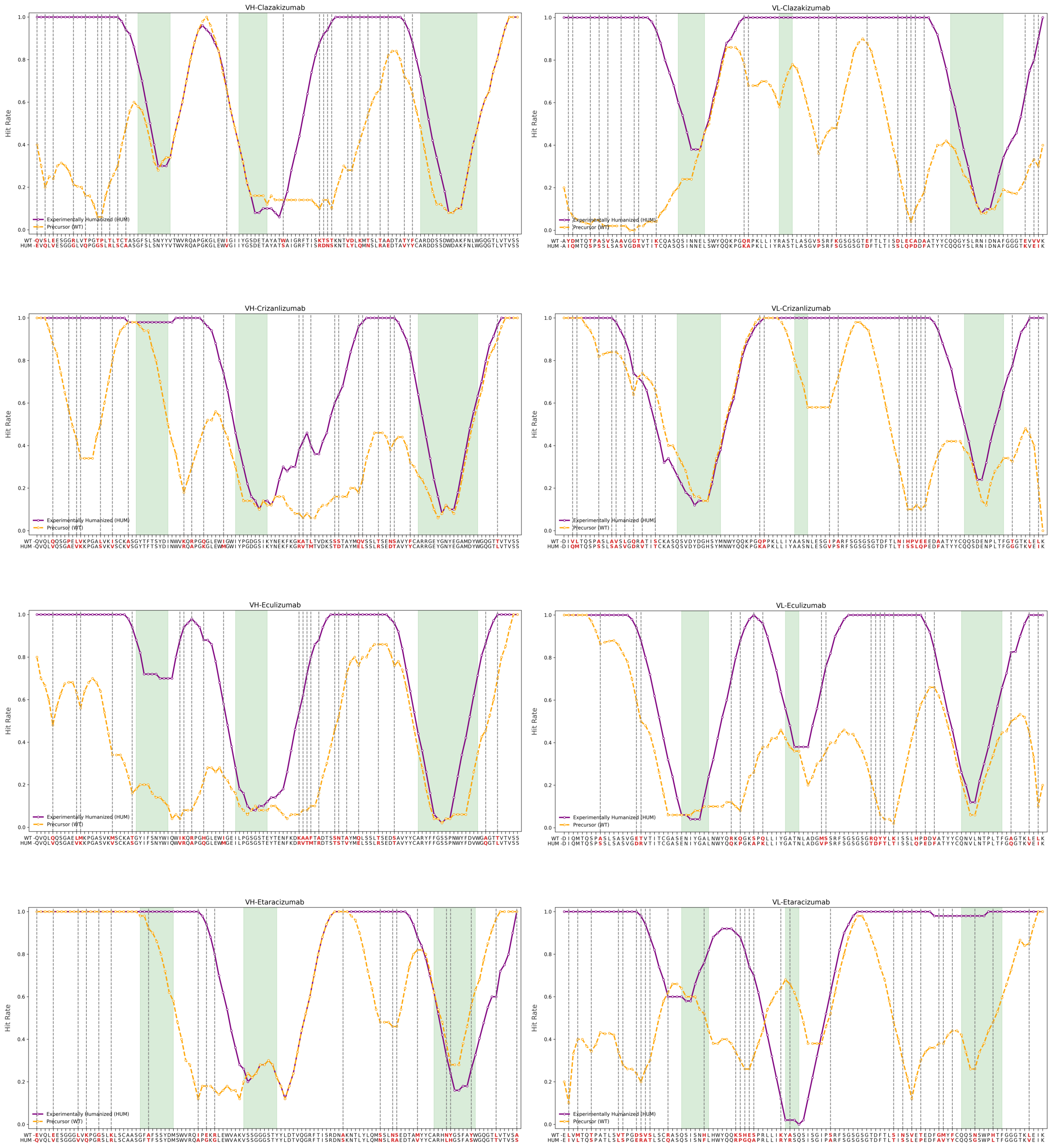


**Supplementary Fig. 4. Site-specific hit rate profiles for heavy chain (VH, left) and light chain (VL, right) of therapeutics antibodies.** From top to bottom: Clazakizumab, Crizanlizumab, Eculizumab, and Etaracizumab. CDR regions are shaded green; mutation sites are marked red. Yellow line: WT hit rate; purple line: humanized hit rate. Humanization mutations preferentially increase hit rates in FRs, with minimal impact on CDRs.

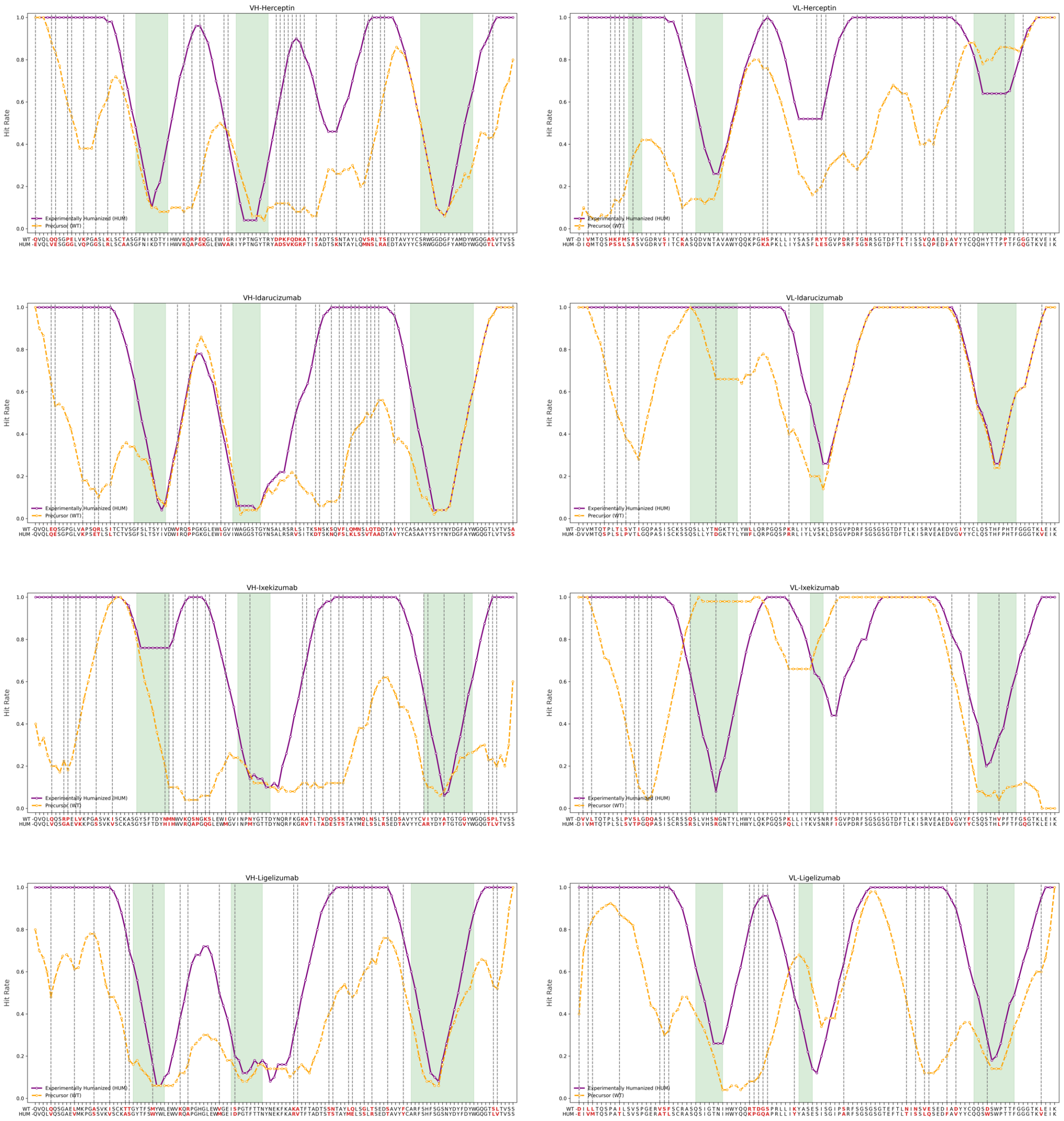


**Supplementary Fig. 5. Site-specific hit rate profiles for heavy chain (VH, left) and light chain (VL, right) of therapeutics antibodies.** From top to bottom: Herceptin, Idarucizumab, Ixekizumab, and Ligelizumab. CDR regions are shaded green; mutation sites are marked red. Yellow line: WT hit rate; purple line: humanized hit rate. Humanization mutations preferentially increase hit rates in FRs, with minimal impact on CDRs.

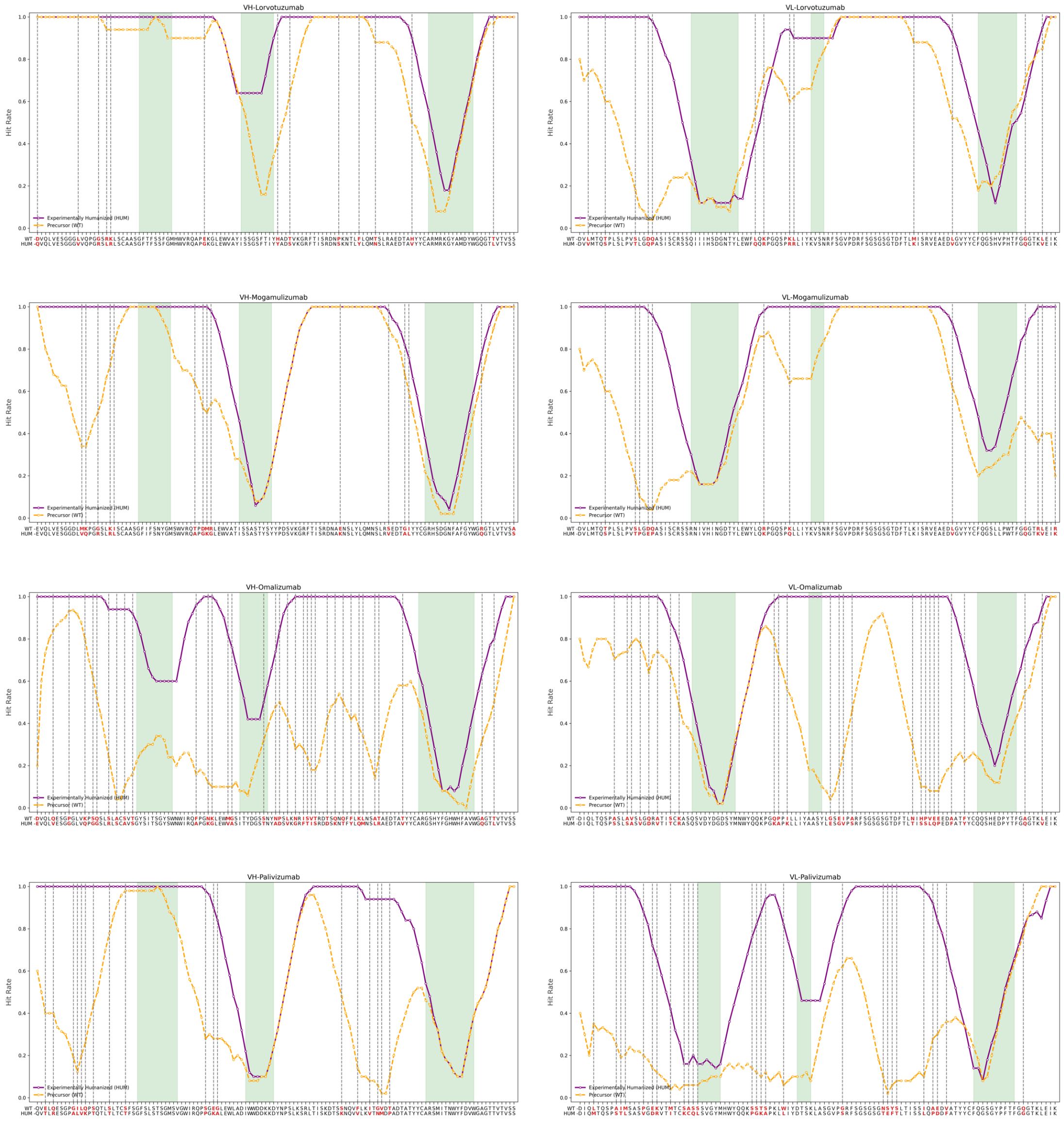


**Supplementary Fig. 6. Site-specific hit rate profiles for heavy chain (VH, left) and light chain (VL, right) of therapeutics antibodies.** From top to bottom: Lorvotuzumab, Mogamulizumab, Omalizumab, and Palivizumab. CDR regions are shaded green; mutation sites are marked red. Yellow line: WT hit rate; purple line: humanized hit rate. Humanization mutations preferentially increase hit rates in FRs, with minimal impact on CDRs.

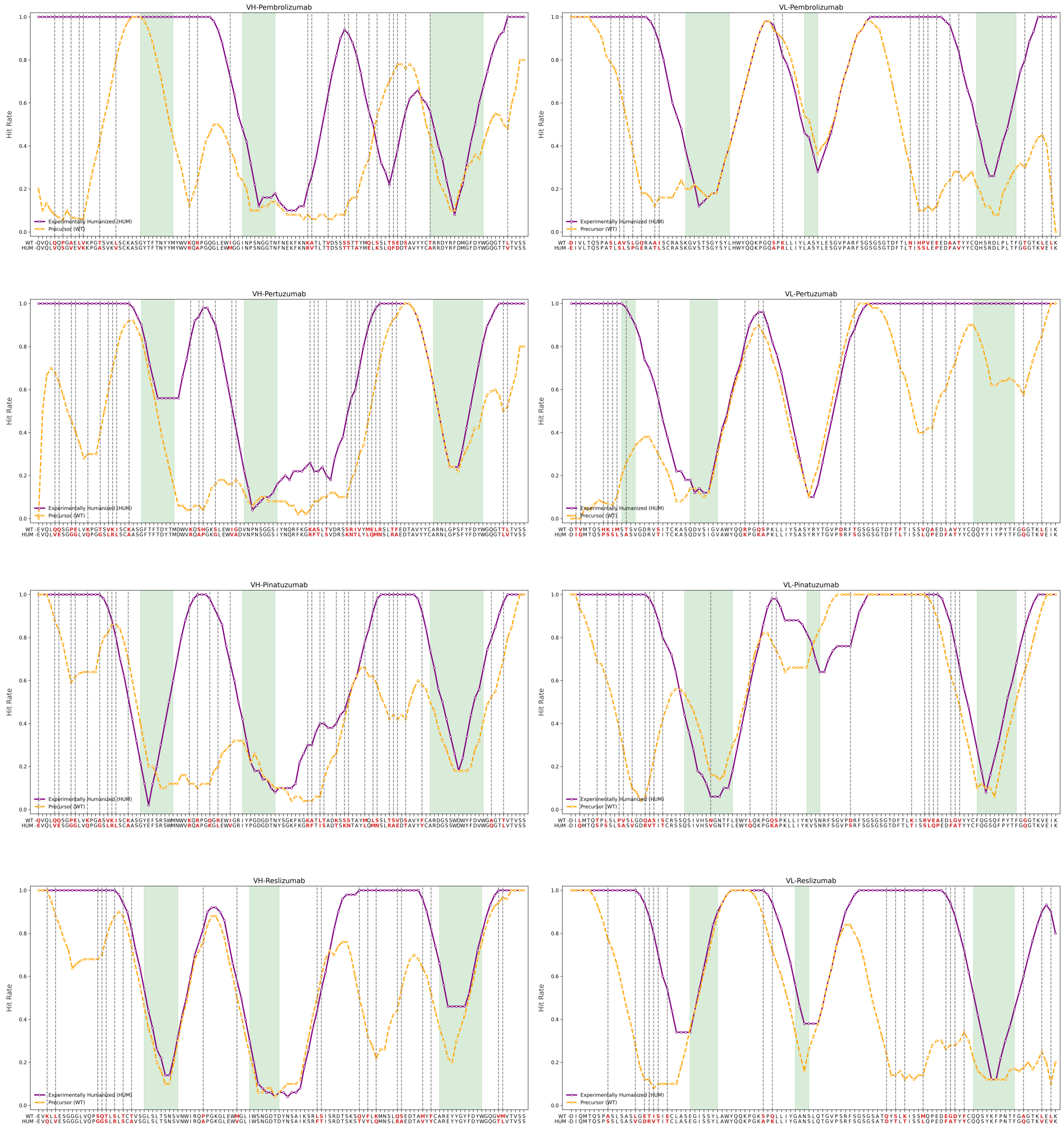

**Supplementary Fig. 7. Site-specific hit rate profiles for heavy chain (VH, left) and light chain (VL, right) of therapeutics antibodies.** From top to bottom: Pembrolizumab, Pertuzumab, Pinatuzumab, and Reslizumab. CDR regions are shaded green; mutation sites are marked red. Yellow line: WT hit rate; purple line: humanized hit rate. Humanization mutations preferentially increase hit rates in FRs, with minimal impact on CDRs.

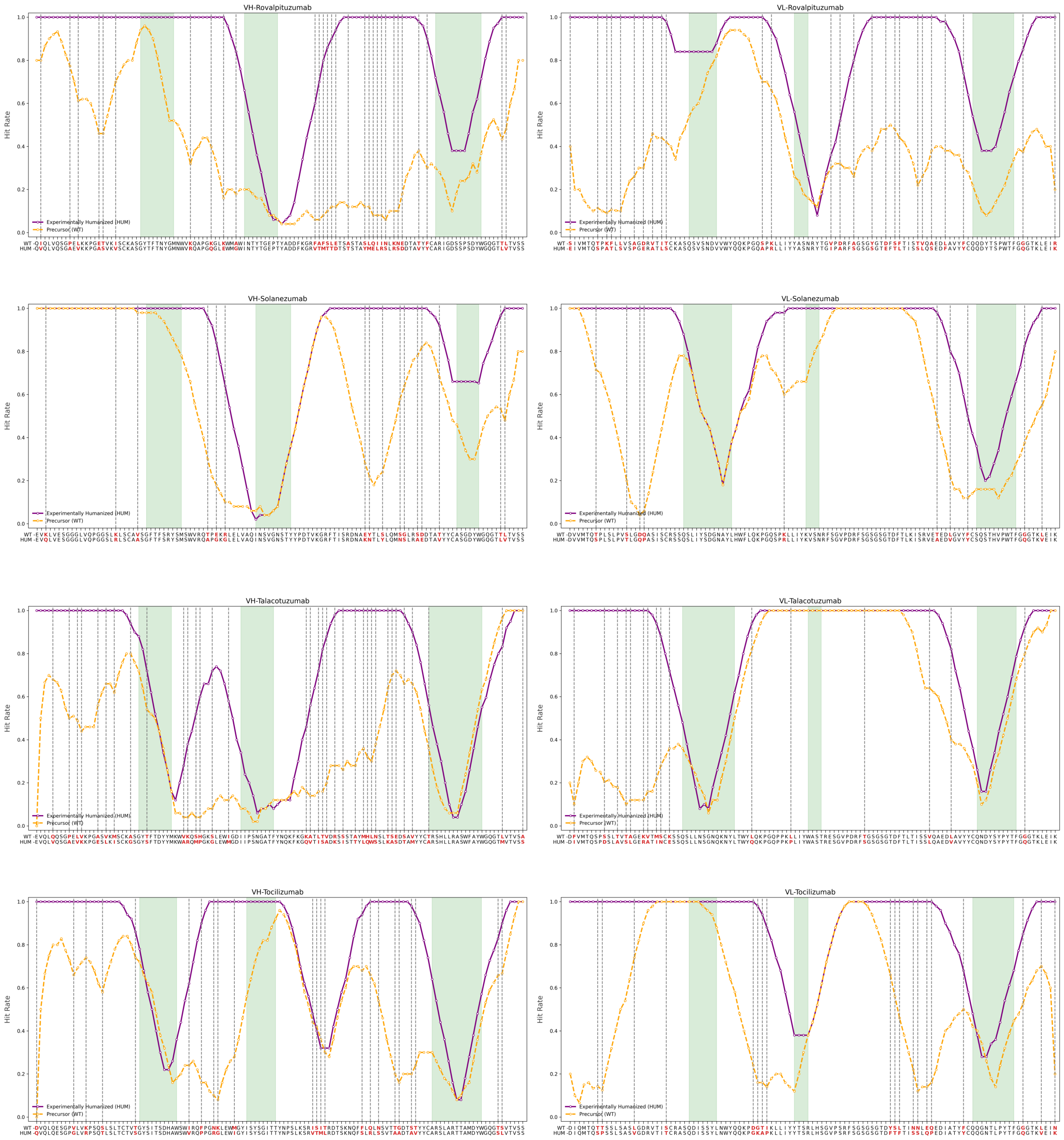


**Supplementary Fig. 8. Site-specific hit rate profiles for heavy chain (VH, left) and light chain (VL, right) of therapeutics antibodies.** From top to bottom: Rovalpituzumab, Solanezumab, Talacotuzumab, and Tocilizumab. CDR regions are shaded green; mutation sites are marked red. Yellow line: WT hit rate; purple line: humanized hit rate. Humanization mutations preferentially increase hit rates in FRs, with minimal impact on CDRs.

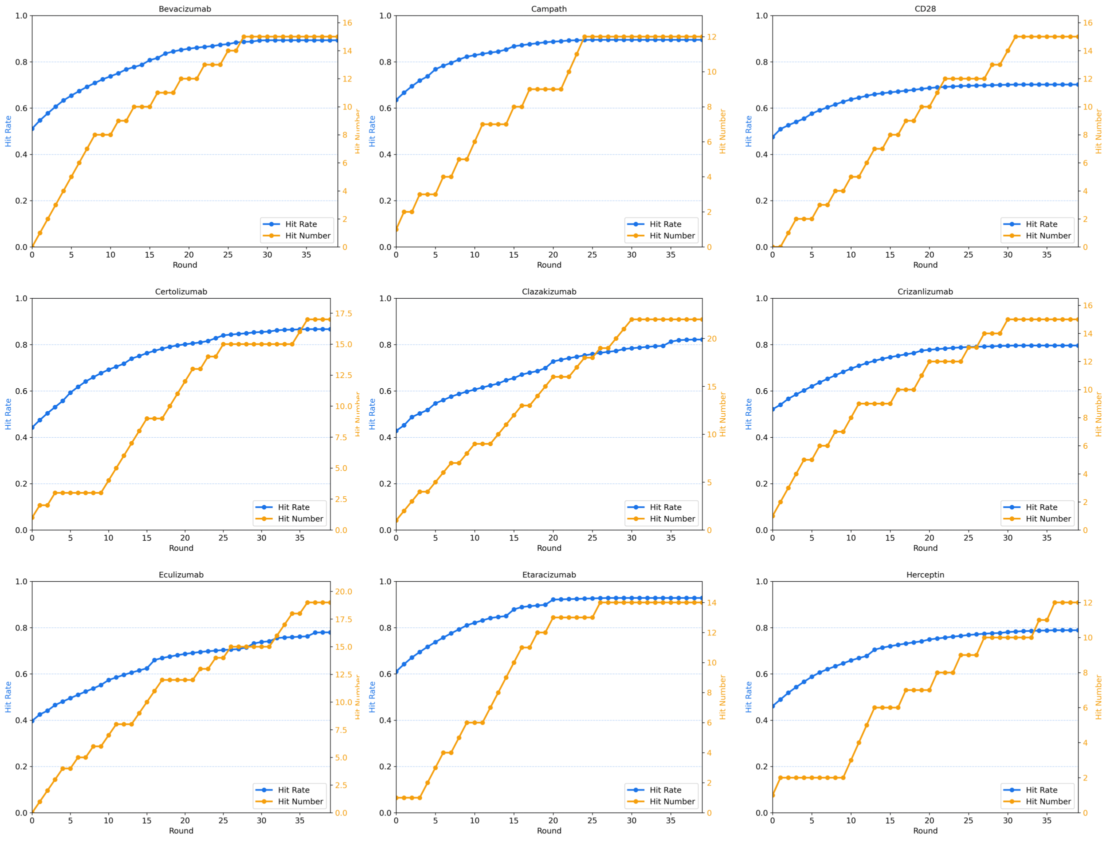


**Supplementary Fig. 9. Humanization design using the greedy iterative strategy.** Detailed iterative optimization of nine antibodies from 25 humanization cases. Hit rate (blue) and hit number (orange) curves demonstrate steady improvement over iterations, converging at 25-30 rounds.

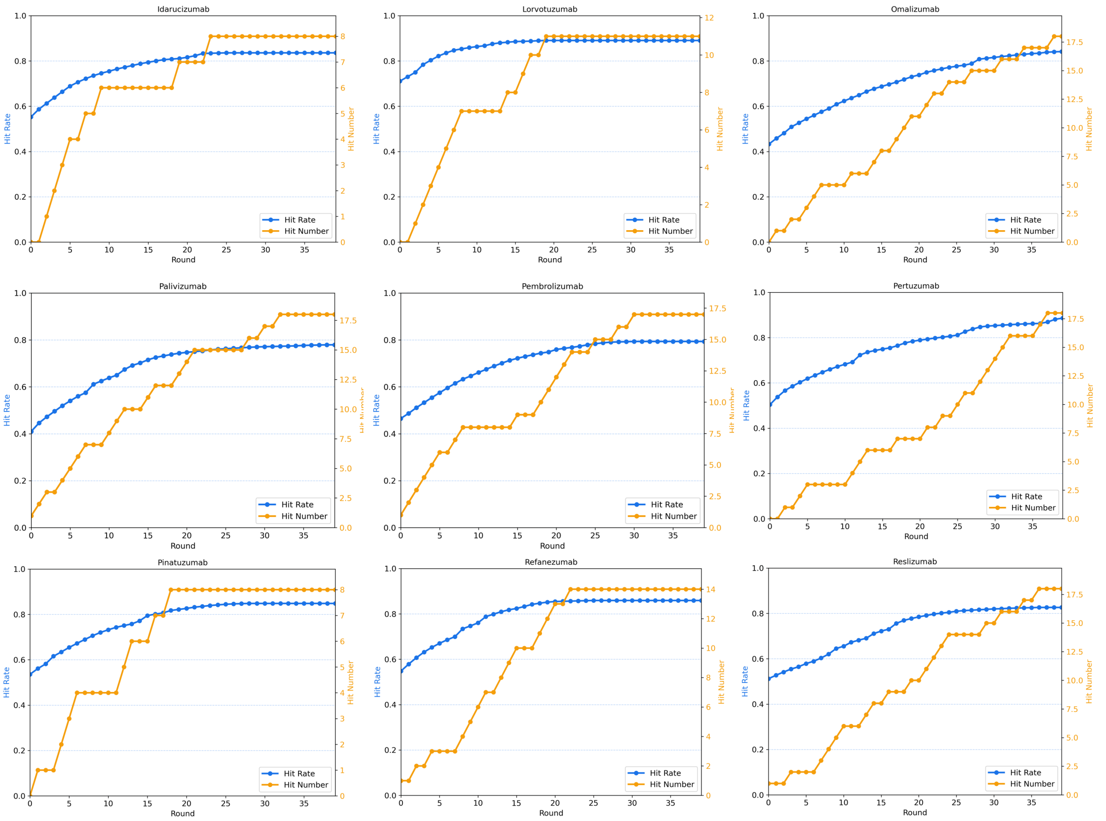


**Supplementary Fig. 10. Humanization design using the greedy iterative strategy.** Detailed iterative optimization of nine antibodies from 25 humanization cases. Hit rate (blue) and hit number (orange) curves demonstrate steady improvement over iterations, converging at 25-30 rounds.

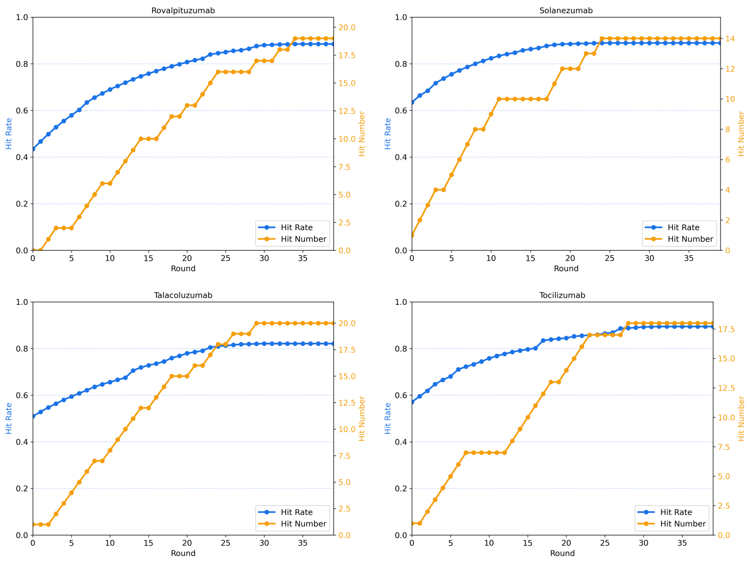


**Supplementary Fig. 11. Humanization design using the greedy iterative strategy.** Detailed iterative optimization of four antibodies from 25 humanization cases. Hit rate (blue) and hit number (orange) curves demonstrate steady improvement over iterations, converging at 25-30 rounds.

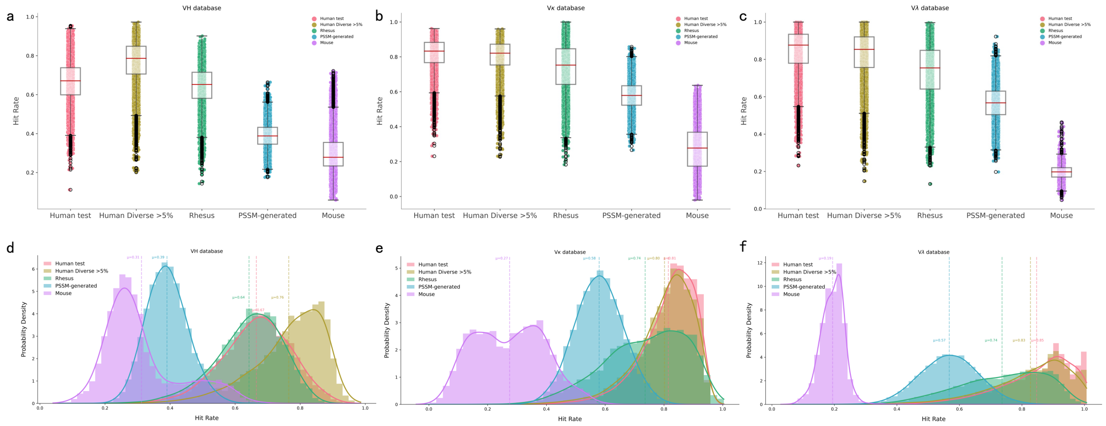


**Supplementary Fig. 12. ImmunoSeq performance in humanness classification. (a-c)** Box plots of hit rate distribution by species in heavy chain database **(a)**, kappa light chain database **(b)**, and lambda light chain database **(c)** from AbNatiV. Hit rate distributions of antibodies from different species (including Human test, Human Diverse (>5%), Rhesus, Mouse, etc.) are shown; boxes represent interquartile ranges (IQRs), horizontal lines denote medians, whiskers extend to 1.5×IQR, and points indicate outliers. **(d-f)** Bar plots of hit rate population by species in heavy chain database **(d)**, kappa light chain database **(e)**, and lambda light chain database **(f)** from AbNatiV. Bar heights represent the relative population of antibodies within each species category, with hit rate on the x-axis and species-colored bars reflecting the distribution of hit rates across species.

**Supplementary Table 1. Evaluation of the ROC-AUC classification task for human heavy chain sequences from AbNatiV**

| Method | Rhesus vs | | PSSM-generated vs | | Mouse vs | |
|---|---|---|---|---|---|---|
| | T | D | T | D | T | D |
| AbNatiV | **0.958** | **0.911** | **0.999** | **0.997** | **0.995** | 0.986 |
| OASis (relaxed) | 0.526 | 0.79 | 0.976 | 0.989 | 0.918 | 0.965 |
| Sapiens | 0.581 | 0.842 | 0.991 | 0.996 | 0.98 | **0.992** |
| ImmunoSeq | 0.558 | 0.807 | 0.981 | 0.991 | 0.976 | 0.991 |

**Supplementary Table 2. Evaluation of the ROC-AUC classification task for human lambda light chain sequences from AbNatiV**

| Method | Rhesus vs | | PSSM-generated vs | | Mouse vs | |
|---|---|---|---|---|---|---|
| | T | D | T | D | T | D |
| AbNatiV | **0.855** | 0.797 | **0.986** | **0.972** | **1** | **1** |
| OASis (relaxed) | 0.791 | 0.792 | 0.943 | 0.936 | 0.999 | 0.998 |
| Sapiens | 0.843 | **0.834** | 0.969 | 0.949 | 0.997 | 0.997 |
| ImmunoSeq | 0.737 | 0.7 | 0.953 | 0.936 | **1** | **1** |

**Supplementary Table 3. Evaluation of the ROC-AUC classification task for human kappa light chain sequences from AbNatiV**

| Method | Rhesus vs | | PSSM-generated vs | | Mouse vs | |
|---|---|---|---|---|---|---|
| | T | D | T | D | T | D |
| AbNatiV | 0.8 | 0.763 | **0.99** | 0.984 | 0.997 | 0.996 |
| OASis (relaxed) | 0.705 | 0.719 | 0.948 | 0.949 | 0.992 | 0.991 |

| | | | | | | |
|---|---|---|---|---|---|---|
| Sapiens | **0.815** | **0.825** | 0.986 | **0.986** | 0.992 | 0.992 |
| ImmunoSeq | 0.672 | 0.642 | 0.961 | 0.946 | **0.999** | **0.998** |

**Supplementary Table 4. Evaluation of the PR-AUC classification task for human heavy chain sequences from AbNatiV**

| Method | Rhesus vs | | PSSM-generated vs | | Mouse vs | |
|---|---|---|---|---|---|---|
| | T | D | T | D | T | D |
| AbNatiV | **0.965** | **0.923** | **1** | **0.998** | **0.996** | 0.988 |
| OASis (relaxed) | 0.57 | 0.829 | 0.982 | 0.992 | 0.897 | 0.965 |
| Sapiens | 0.626 | 0.883 | 0.993 | 0.997 | 0.982 | **0.994** |
| ImmunoSeq | 0.578 | 0.842 | 0.985 | 0.994 | 0.976 | 0.992 |

**Supplementary Table 5. Evaluation of the PR-AUC classification task for human lambda light chain sequences from AbNatiV**

| Method | Rhesus vs | | PSSM-generated vs | | Mouse vs | |
|---|---|---|---|---|---|---|
| | T | D | T | D | T | D |
| AbNatiV | 0.809 | 0.769 | **0.992** | 0.988 | 0.998 | **0.997** |
| OASis (relaxed) | 0.734 | 0.744 | 0.959 | 0.961 | 0.993 | 0.993 |
| Sapiens | **0.848** | **0.86** | 0.989 | **0.989** | 0.993 | 0.993 |
| ImmunoSeq | 0.597 | 0.572 | 0.97 | 0.96 | **0.999** | **0.997** |

**Supplementary Table 6. Evaluation of the PR-AUC classification task for human kappa light chain sequences from AbNatiV**

| Method | Rhesus vs | PSSM-generated vs | Mouse vs |
|---|---|---|---|

| | T | D | T | D | T | D |
|---|---|---|---|---|---|---|
| AbNatiV | 0.861 | 0.805 | **0.99** | **0.98** | **1** | **1** |
| OASis (relaxed) | 0.818 | 0.822 | 0.958 | 0.955 | **1** | 0.999 |
| Sapiens | **0.876** | **0.877** | 0.978 | 0.967 | 0.999 | 0.999 |
| ImmunoSeq | 0.756 | 0.713 | 0.965 | 0.953 | **1** | **1** |

**Supplementary Table 7. Hit rate of the humanization design sequences**

| | Experiment | Hu-mAb | Sapiens | ImmunoSeq |
|---|---|---|---|---|
| Average hit rate | 0.756 | 0.619 | 0.760 | **0.834** |
| Average mutation number | 45.08 | 25.32 | 32.72 | 32.38 |
